# Photoinitiator-free visible-light bioprinting: minimising radical-induced oxidative stress and DNA damage in gelatin hydrogels

**DOI:** 10.64898/2025.12.01.691472

**Authors:** Samaneh Eftekhari, Xin Y. Oh, Daren Zhou, Jessica E. Frith, Helena C. Parkington, John S. Forsythe, Vinh X. Truong, Timothy F. Scott

**Affiliations:** Department of Materials Science and Engineering, Monash University, Clayton, Victoria, 3800 Australia; Institute of Sustainability for Chemicals, Energy and Environment (ISCE2), Agency for Science, Technology and Research (A*STAR), 1 Pesek Road, J’urong Island, Singapore, 627833 Singapore; ARC Training Centre for Cell and Tissue Engineering Technologies, Monash University, Clayton, Victoria, 3800 Australia; Department of Physiology, Biomedicine Discovery Institute, Monash University, Clayton, VIC, Australia; Department of Chemical and Biological Engineering, Monash University, Clayton 3800, Australia

**Keywords:** Photocrosslinkable hydrogel, Initiator-free photopolymerization, Visible light crosslinking, 3D bioprinting, 3D modelling

## Abstract

Light-mediated crosslinking of polymers is widely employed in the preparation of hydrogels for biofabrication and tissue engineering, since photo-crosslinking enables the spatiotemporal control over the gelation processes. Nevertheless, a critical bottleneck persists: most photo-crosslinking reactions rely on the use of photo-initiator, and ultraviolet or short-visible-wavelength light activation, which suffers from potential photodamage and poor penetration. Here we present a photoinitiator-free hydrogel system based on gelatin functionalized with acrylamidylpyrene groups (Gel-Pyr) able to undergo crosslinking *via* visible-light-induced [2+2] cycloaddition. Gel-Pyr solution exhibits rapid gelation kinetics, tuneable mechanical properties, facile temporal control over photocrosslinking, and long-term structural stability (>30 days) in cell culture conditions. Rheological analyses reveal pronounced shear-thinning behaviour at room temperature, enabling extrusion-based 3D bioprinting of multilayered constructs with high structural fidelity. Fine strand resolution (<400 µm) is achieved in bioprinted crosshatch structures, enabling sufficient nutrient diffusion for cell support. Compared with gelatin methacryloyl (GelMA), Gel-Pyr significantly reduces photocrosslinking-induced oxidative stress and apoptosis in encapsulated bone-marrow mesenchymal stem cells (BM-MSCs), supporting >80% viability over 7 days. By eliminating UV exposure and lowering free radical generation, this visible-light-responsive hydrogel platform offers a facile and cytoprotective alternative to other hydrogel systems.

**Table of Content:** A visible light-crosslinkable, initiator-free gelatin-based hydrogel (Gel-Pyr) is developed using acrylamidylpyrene functionalization. This system enables rapid crosslinking under cytocompatible reaction conditions and offers excellent printability, tuneable mechanics, and long-term stability. Gel-Pyr supports high cell viability, reduced oxidative stress and precision bioprinting, positioning it as a promising platform for tissue engineering and *in vitro* biofabrication.

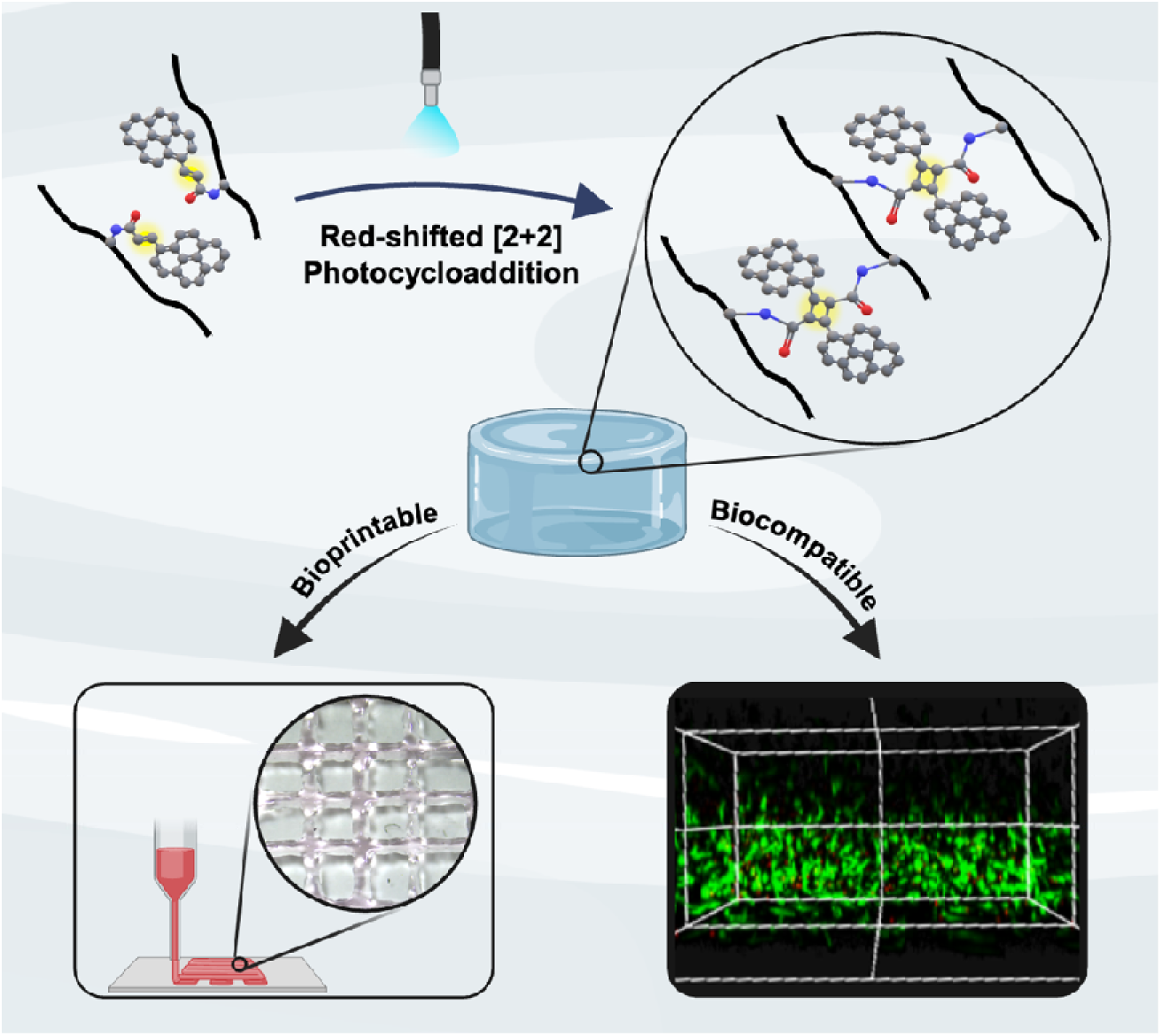

## 1 Introduction

Creating three-dimensional hydrogel systems that mimic natural extracellular matrix (ECM) requires materials that are biocompatible, chemically and mechanically tuneable, and capable of supporting cellular function. Due to their similarity to native ECM in both structure and function, hydrogels offer tremendous potential in tissue engineering and regenerative medicine applications and the development of tissue specific *in vitro* models (1–4), where they support cell adhesion (5), proliferation (6), and differentiation (7). In addition, hydrogels offer the potential for spatiotemporal control over gelation kinetics and microarchitecture through external stimuli such as light, enabling precise patterning of cell-laden structures (8).

Methacryoyl functionalised systems such as gelatin methacryoyl (GelMA) (9), hyaluronic acid methacryoyl (HAMA) (10) and alginate methacryoyl (ALMA) (11) are widely used to afford mechanically-stable hydrogels (1, 12, 13). Photopolymerization is commonly employed for chemically crosslinking these hydrogels, owing to its spatiotemporal reaction control and compatibility with physiological reaction conditions. This method generally relies on water-soluble photoinitiators, such as Irgacure 2959 and lithium phenyl-2,4,6-trimethylbenzoylphosphinate (LAP), which are activated by (ultra)violet light to trigger covalent crosslinking *via* a free-radical polymerization process (12, 14). Despite their widespread use, these photoinitiators present notable limitations. Irgacure 2959, for example, has a low molar extinction coefficient (∼4 M ¹·cm ¹) at 365 nm, necessitating either prolonged UV exposure, increased light intensity, or comparably higher initiator concentrations to achieve sufficient crosslinking, conditions that have been shown to impair cell viability due to cytotoxic effects (15). Similarly, while LAP is often considered more biocompatible (with about 96% cell survival with 2.2 mM concentration), it still presents cytotoxicity concerns at higher concentrations (15, 16). In addition to photoinitiator toxicity, the reactive free radicals generated during photopolymerization can damage cellular components, disrupt membranes, and induce oxidative stress, apoptosis, or senescence, thereby compromising encapsulated cell function and viability (17, 18). UV light itself can contribute to oxidative stress, mutagenicity and phototoxicity (19, 20). These adverse effects can also compromise the functional performance of engineered tissues. For example, in 3D-bioprinted liver constructs, excess free radicals generated during GelMA photocrosslinking impaired hepatocyte-specific functions, including albumin secretion and urea production, necessitating the incorporation of radical scavengers such as ascorbic acid and α-tocopherol to restore tissue-level functionality (21, 22). Likewise, increasing photoinitiator concentrations during GelMA crosslinking has been shown to elevate intracellular ROS levels in mesenchymal stromal cells, resulting in reduced viability, disrupted focal adhesions, and impaired cell spreading (23). Collectively, these studies underscore the need for hydrogel systems that preserve the processing advantages of photocrosslinking, including rapid gelation, precise spatial control, and robust mechanical properties, while overcoming the cytotoxic limitations that continue to hinder the broader application of cell-laden hydrogels in advanced biofabrication. Here, we introduce a hydrogel system that enables rapid, catalyst-free crosslinking under visible light irradiation, minimizing free radical generation. Based on an acrylamidylpyrene chromophore, this system is synthesised by conjugating pyrene acrylic acid to the amino groups of gelatin, yielding a chromophore-functionalized gelatin referred to here as gelatin-pyrene (Gel-Pyr). Upon irradiation with visible light (400–500 nm), Gel-Pyr undergoes [2+2] photocycloaddition to generate a cyclobutane adduct, enabling rapid and additive-free hydrogel cross-linking. The absence of photoinitiators simplifies hydrogel preparation, enhances its usability as a bioink and, importantly, eliminates free radical generation. We further demonstrate the tuneable mechanical properties, bioprintability, and cytocompatibility of the Gel-Pyr hydrogel system, highlighting its potential as an alternative to contemporary hydrogel systems such as Gel-MA and enabling advanced tissue engineering and 3D modelling applications.

## 2 Results and Discussion

### 2.1 Photoinitiator-free, visible light gelatin-pyrene photo-crosslinking

Gelatin-acrylamidylpyrene (Gel-Pyr) was synthesized by reacting a succinimidyl ester of pyrene acrylic acid with amino groups borne by the lysine residues of gelatin **(Figure 1a)**. Unlike our previous work on [2+2] photocycloaddition of styrylpyrene functionalised polyethylene glycol (PEG) hydrogels(24), acrylamidylpyrene functionalisation of gelatin (Gel-Pyr) afforded improved solubility in aqueous environments. Given the intended application for cell encapsulation, Dulbecco’s Modified Eagle Medium (DMEM) was chosen as the solvent for characterisation of the hydrogel properties. Ultraviolet-visible (UV-vis) spectroscopic analysis of Gel-Pyr in DMEM revealed a distinct absorption peak at 369 nm, accompanied by a shoulder around 405 nm (**Figure 1b**). In contrast, the absorbance spectrum of pure gelatin showed no absorbance in this region, confirming that the peak was attributable to acrylamidylpyrene functionalization. Thus, to assess the photodimerization extent of the Gel-Pyr-borne acrylamidylpyrene groups, the absorbance spectrum was recorded after visible light exposure, where the hydrogel precursor solution was irradiated with 405 nm light at an intensity of 20 mW.cm^-2^ for 2 minutes. Upon irradiation, absorbance peaks at 335 and 352 nm emerged, reminiscent of the pyrene absorbance spectrum (25, 26), while the 369 nm peak was eliminated, indicating near-quantitative consumption of the acrylamidylpyrene groups *via* [2+2] photocycloaddition to afford the cyclobutane adduct bearing pyrene pendant groups. Subsequently, irradiation of a Gel-Pyr solution to 405 nm light for 3 minutes yielded a clear hydrogel **(Figure 1c)**, confirming both the successful functionalization of gelatin with the acrylamidylpyrene moiety, as well as the visible light-mediated cross-linking of Gel-Pyr *via* a photocycloaddition reaction.

**Figure 1.**
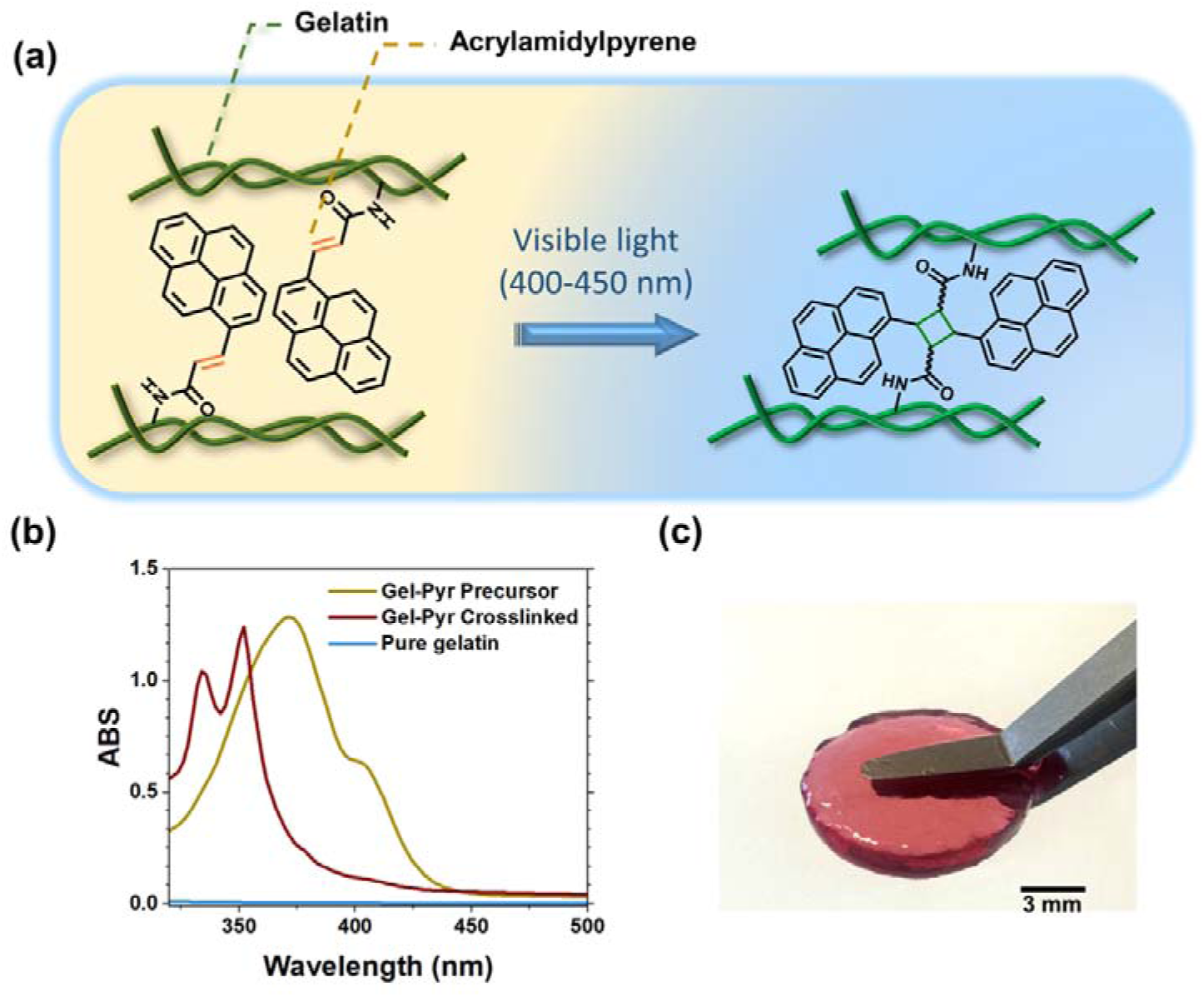
Synthesis and visible-light cross-linking of Gel-Pyr hydrogels. a) Scheme of polymer photo-crosslinking *via* photocycloaddition of acrylamidylpyrene. b) UV-vis spectra of gelatin-pyrene in DMEM culture medium before (yellow line) and after (red line) 405 nm light irradiation; followed by a comparison with pure gelatin in cell culture medium (blue line). c) Gel-Pyr 7.5% w/v cross-linked in DMEM under 405 nm light irradiation for 3 min. The scale bar represents 3 mm.

The degree of functionalization, representing the percentage of lysine groups reacted with acrylamidylpyrene-NHS, was estimated using a calibration curve of acrylamidylpyrene- amide in dimethyl sulfoxide (DMSO) (**Figure S1, Supporting Information**). The concentration of acrylamidylpyrene groups borne by the functionalized gelatin (i.e., Gel-Pyr) at 0.0625% w/v in DMSO was calculated to be 5.51 × 10^-5^ M. Considering that porcine skin gelatin contains approximately 2.7% lysine residues by weight (27, 28), the degree of functionalization (DoF) of lysine groups was determined to be 47%. The DoF is a critical parameter that directly governs crosslinking density and network stiffness, higher DoF producing stiffer matrixes, whilst lower DoF preserves free lysine residues and the associated cell-instructive bioactivity of the gelatin backbone (29, 30). The intermediate DoF achieved here, provides a balance between efficient network formation and retention of the biological functionality of gelatin, enabling the formation of mechanically robust hydrogels while maintaining its inherent bioactivity.

### 2.2 Gel-Pyr is a biomechanically tuneable hydrogel with viscoelastic and shear-thinning properties

A comprehensive understanding of the rheological properties of hydrogels is critical for establishing the relationship between their chemical composition and macroscopic behaviour. This knowledge is fundamental to designing hydrogels with tailored properties for specific applications, including tissue engineering, drug delivery, and 3D printing (31). In tissue engineering, for instance, hydrogels serve as scaffolds that must replicate the mechanical properties of native tissues, as these properties directly affect cell behaviour, mechanotransduction, and overall cellular function (32, 33). Furthermore, the use of hydrogels as inks for bioprinting requires a specific suite of physical properties to enable fabrication of precisely defined structures.

Accordingly, a comprehensive analysis was conducted to interrogate the rheological properties of Gel-Pyr, focussing on the properties both prior to light exposure and after photo-crosslinking. The dynamic properties of the Gel-Pyr solutions ranging from 3.5% to 10.0% w/v was evaluated at 25°C prior to light exposure across applied frequency and strain ranges to assess their viscoelastic performance under varying mechanical conditions. The *G*′ of Gel-Pyr exceeded the *G*″ at all concentrations examined in the dynamic oscillation frequency scan, indicating predominantly elastic behaviour **(Figure 2a)**. Furthermore, the strain sweep results **(Figure 2b)** showed that the *G*′ plateaued across the entire strain range for all concentrations, signifying an extended linear viscoelastic region. These data suggest the formation of solid-gel structure in which the coordinated physical cross-linked network effectively resists mechanical disruption throughout the tested range. The chemical gelation kinetics of the hydrogel were further determined using *in situ* photo-rheometry measurements, enabling real-time monitoring of the hydrogel’s mechanical evolution during light-induced crosslinking. **Figure 2c** demonstrates *G*′ for all concentrations examined at 25°C. The onset of light irradiation triggered a rapid photo-crosslinking reaction, where *G*′ increased sharply following 1 s light irradiation, with complete gelation (90% of plateau) achieved within 43 - 90 s; specifically, *G*′ increased from 4 ± 1 to 58 ±7 Pa for 3.5% w/v, from 9 ± 2 to 377 ± 118 Pa for 5.0% w/v, from 23 ± 2 Pa to 1.5 ± 0.1 kPa for 7.5% w/v, and from 34.0 ± 0.2 Pa to 3.0 ± 0.2 kPa for 10% w/v. Significant differences in *G*′ were evident among all concentration groups, except between 3.5% and 5.0% w/v **(Figure 2d)**. As expected, higher concentrations resulted in increased *G*′, reflecting greater hydrogel stiffness. Notably, a slight increase in *G*′ was observed in the 7.5% and 10.0% w/v Gel-Pyr groups prior to light irradiation, due to enhanced chain entanglement and possible association of the pyrene groups. The compressive properties of the bulk hydrogel were also assessed using compression testing **(Figure 2e)**. The compressive (Young’s) modulus, a key parameter in determining hydrogel elasticity, plays a crucial role in 3D modelling and tissue engineering applications, where balancing mechanical integrity and biocompatibility is essential (34). Testing of 3.5%, 5.0% and 7.5% w/v Gel-Py hydrogels showed that 7.5% Gel-Pyr exhibited the highest Young’s modulus at 12 ± 1 kPa, followed by values of 4.2 ± 0.8 kPa and 2.8 ± 0.1 kPa for 5.0% and 3.5% w/v hydrogels, respectively, though the difference between these two was not statistically significant. This variation in Young’s modulus across different concentrations can be attributed to the differences in the volume fraction of gelatin and crosslink density, both of which contribute to enhanced network rigidity and increased polymer chain interactions at raised gelatin concentrations. Based on these findings, the 7.5% w/v Gel-Pyr composition was selected for subsequent characterization, as its stiffness fell within a range considered suitable for the intended cellular applications.

**Figure 2.**
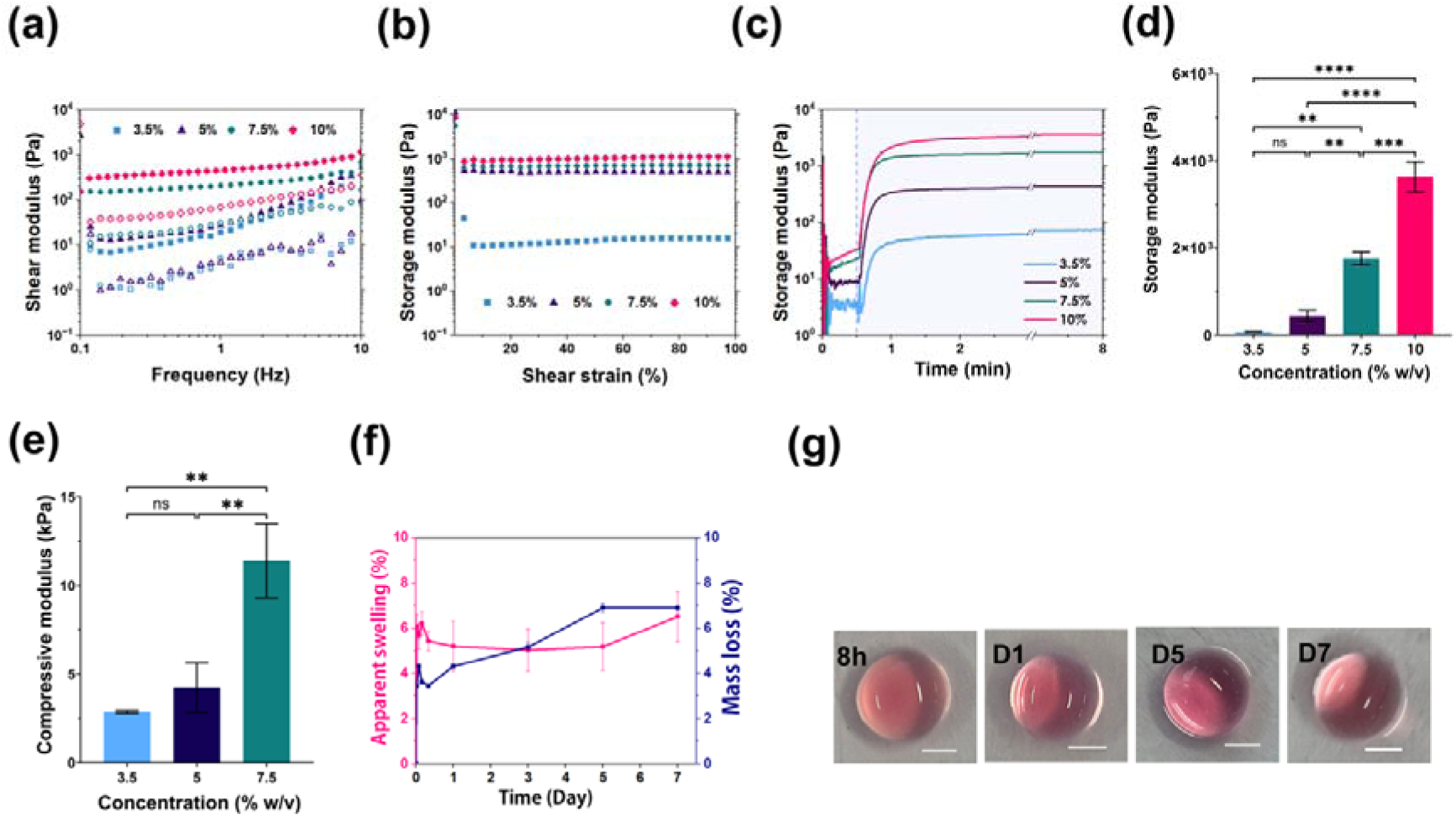
Mechanical characterization and swelling behaviour of Gel-Pyr hydrogels. (a) Shear modulus, *G*′ (closed symbol) and *G*″ (open symbol), of Gel-Pyr at concentrations of 3.5% w/v, 5.0% w/v, 7.5% w/v, and 10.0% w/v at 25°C as a function of frequency at a fixed strain (1%). (b) Storage modulus of Gel-Pyr (3.5% w/v, 5.0% w/v, 7.5% w/v, and 10.0% w/v) at 25°C as a function of strain at a fixed frequency (1 Hz). (c) Photo-crosslinking evolution for Gel-Pyr at concentrations of 3.5% w/v, 5.0% w/v, 7.5% w/v, and 10.0% w/v at 25°C using visible light irradiation with the intensity of 20 mW·cm^-^², demonstrating a rapid increase in storage modulus (*G*′) upon irradiation. Light irradiation was initiated after a 30 s equilibration period indicated by a dashed line. (d) Maximum storage modulus (*G*′) achieved at each concentration (n = 3, ****P < 0.0001, mean ± S.E.M., one-way ANOVA). (e) Young’s modulus of chemically cross-linked Gel-Pyr measured at 10-20% compressive strain, comparing different concentrations after equilibrium swelling (n = 3, **P < 0.01, mean ± S.E.M., one-way ANOVA). (f) Apparent swelling and mass loss of Gel-Pyr (7.5% w/v) in DMEM culture medium to assess the swelling kinetics of Gel-Pyr over 7 days (n = 3, mean ± S.E.M.). (g) Representative images of swollen 7.5% w/v Gel-Pyr hydrogel at 8h, days 1, 5 and 7. Scale bar: 2.5 mm.

The swelling behaviour of hydrogels is a key parameter that reflects their network structure, crosslink density, and interaction with its surrounding environment (35). Thus, the swelling and degradation profile of photo-crosslinked Gel-Pyr hydrogels were evaluated over a 7-day period under physiologically relevant conditions of 37°C and 5% CO **(Figure 2f)**. The hydrogels demonstrated rapid equilibration during the initial incubation period, reaching an apparent swelling ratio of approximately 5% within the first 8 h, after which the swelling profile remained relatively constant. The limited increase in swelling indicates that the hydrated polymer network remained structurally stable without excessive expansion or network disruption. In parallel, Gel-Pyr hydrogels showed limited mass loss, remaining below 7% after 7 days, with the majority of the loss occurring within the first 24 h. The gradual increase in mass loss over time likely reflects the release of loosely bound polymer chains and unreacted components. Importantly, the comparable progression of swelling and degradation behaviour suggests gradual network relaxation rather than rapid bulk erosion or structural collapse. **Figure 2g** demonstrates the swollen hydrogel over time. Such balanced swelling-degradation kinetics are advantageous for long-term tissue engineering applications, where maintenance of dimensional stability and sustained matrix integrity are essential. Similarly, the crosslinking efficiency of Gel-Pyr network was evaluated by gravimetrically determining the gel fraction of 7.5% w/v hydrogels. The photocrosslinked hydrogel exhibited a gel fraction of 97 ± 0.4%, indicating the formation of a crosslinked network with minimal sol content. This high polymer retention after 48 h of aqueous extraction highlights the efficiency of the light-mediated [2+2] cycloaddition mechanism in forming covalent inter-chain bonds; however physical crosslinking of the Gel-Pyr may also contribute to the gel fraction. Such a high gel fraction is consistent with step growth polymerization processes, where the high functionality of the reacting groups and high conversion rates minimize the amount of extractable fractions ^[21]^. In comparison, conventional free-radical-based systems such as GelMA typically yield gel fractions in the range of 65-80%, depending on the degree of methacrylation and UV exposure conditions ^[22]^.

To evaluate temperature-responsive behaviour of the 7.5% w/v Gel-Pyr formulation, changes in *G*′ and *G*″ were monitored during cooling (40°C to 4°C) and heating (4°C to 40°C) temperature ramps **(Figure 3a)**. Whereas the Gel-Pyr solution exhibited liquid-like behaviour at 40°C, a progressive increase in storage modulus was observed with decreasing temperature, where the *G*′ value increased from 4.2 Pa at 40°C to 7170 Pa at 4°C, while a solution-to-gel transition (*T*_sol-gel_), attributable to physical gelation and marked by a significant increase in *G*′, occurred at 27°C. During the subsequent heating ramp, the modulus remained relatively constant up to nearly 25°C, after which a decline in *G*′ was noted; a gel-to-solution (*T*_gel-sol_) transition observed at 36°C indicating gelation thermo-reversibility. Notably, G′ was higher than *G*″ even at physiological temperatures (*G*′ ∼ 10 Pa), potentially a result of π–π stacking interactions amongst pyrene groups maintaining a degree of elasticity even at elevated temperatures (36, 37). The thermal hysteresis of gelation and degelation can be attributed to the physical cross-linking mechanism where the gelatin backbone undergoes a coil-to-helix transition upon cooling, forming junction zones that function as physical cross-links. Even though these helical structures are stable at low temperatures and dissociate upon heating, they require greater energy input, and consequently higher temperatures, to dissociate and convert the gel into a liquid, as observed in the heating cycle (38–42). Consistent with the observed thermally induced gelation, time-sweep rheology at 20°C revealed progressive network maturation, characterized by a continuous increase in *G*′ to approximately 1 kPa while remaining substantially higher than *G*″, indicative of stable hydrogel formation **(Figure S2, Supplementary Information).**

**Figure 3.**
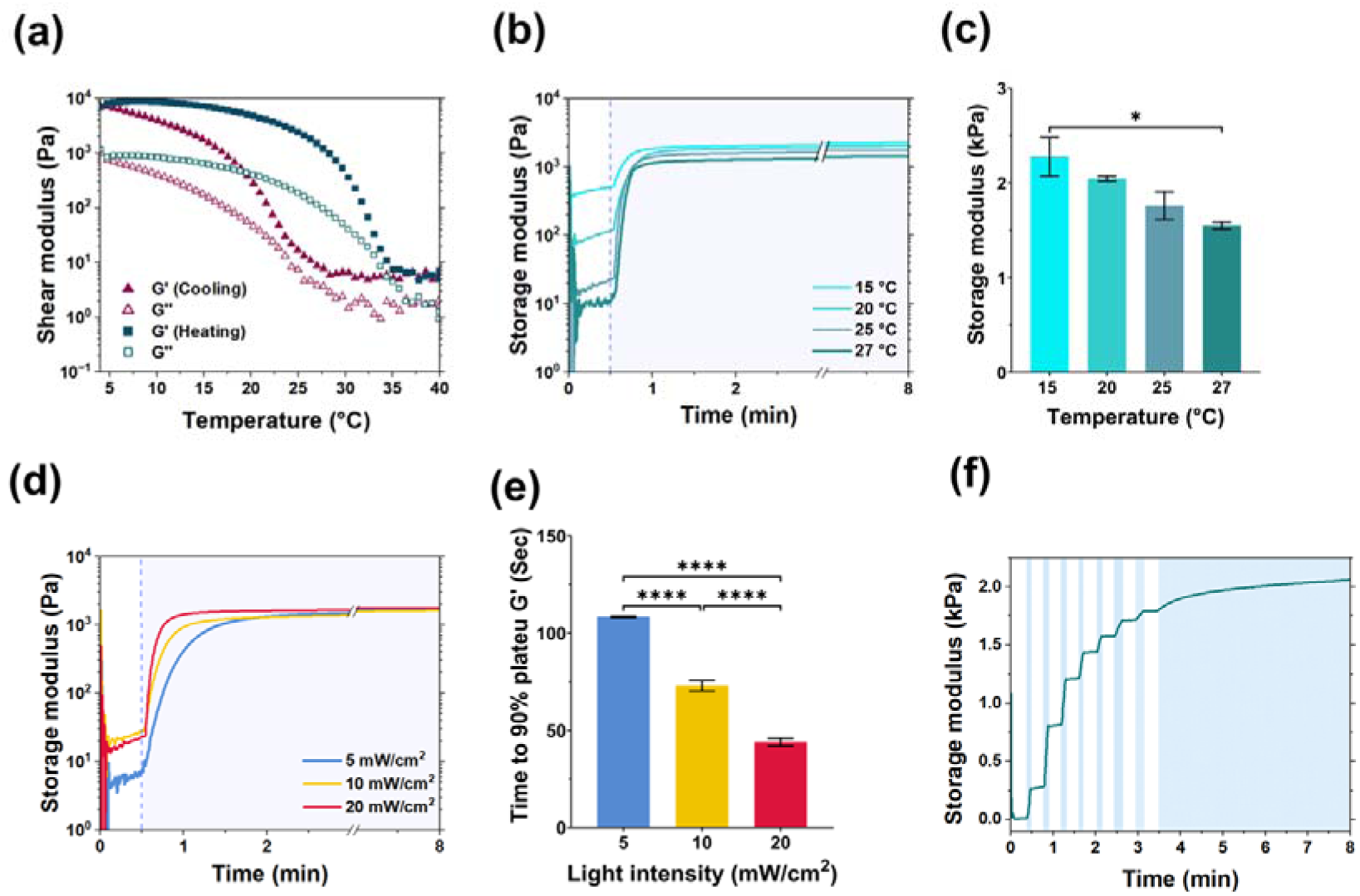
Modulation of Gel-Pyr crosslinking by temperature and light intensity and irradiation duration. (a) Shear modulus as a function of temperature (4°C - 40°C) during a cooling and heating cycle at temperature ramp rates of -1°C·min^-1^ and +1°C·min^-1^, respectively. The graph shows the storage (*G*′) and loss (*G*″) moduli at a constant oscillatory frequency (1 Hz) and applied strain (1%). (b) *G*′ evolution for Gel-Pyr 7.5% w/v at varying temperatures using an irradiation intensity of 20 mW·cm^-^², illustrating the temperature-dependent kinetics of physical crosslinking prior to light exposure. (c) Maximum *G*′ at each temperature (n = 3, *P < 0.05, mean ± S.E.M., one-way ANOVA). (d) *G*′ as a function of light intensity for Gel-Pyr 7.5% w/v at 25°C. (e) Time required to reach 90% of the plateau *G*′ under different light irradiation conditions for Gel-Pyr 7.5% w/v at 25°C (n = 3, ****P < 0.0001, mean ± S.E.M., one-way ANOVA). (f) Time required to reach 90% of the plateau *G*′ under different light irradiation conditions for Gel-Pyr 7.5% w/v at 25°C (n = 3, ****P < 0.0001, mean ± S.E.M., one-way ANOVA). (f) Demonstration of temporal control over crosslinking in Gel-Pyr (7.5% w/v) through alternating period of irradiation (at an intensity of 20 mW·cm^-^²) and darkness. Periodic light exposure intervals are indicated by light blue shading, highlighting the reversible control of network formation in response to irradiation.

The photocrosslinking kinetics of Gel-Pyr over a temperature range of 15-27°C were further investigated across concentrations of 3.5-10% w/v. Representative data for the 7.5% w/v composition are presented in **Figure 3b**, while the corresponding datasets for the remaining concentrations are provided in **Figure S3, Supplementary Information**. Prior to irradiation, the initial *G*′ values were notably higher at 15 and 20°C. This is likely attributed to the formation of triple helices stabilized by hydrogen bonds between gelatin chains at temperatures below the hydrogel’s sol-gel transition temperature (*T*_sol-gel_=27°C) and π-π stacking of the pyrenes. However, the final *G*′ values remained largely consistent across all temperature groups, with only the 15 and 27°C conditions exhibiting statistically significant differences **(Figure 3c)**. The influence of light intensity on the crosslinking kinetics of Gel- Pyr was additionally investigated by employing three blue light intensities: 5, 10 and 20 mW·cm^-2^ **(Figure 3d, 3e)**. The results revealed that, while the maximum *G*′ value remained consistent across all conditions, the time required to reach the plateau varied significantly. The hydrogel achieved a plateau at 44 s, 73 s, and 108 s under light intensities of 20, 10, and 5 mW·cm^-^², respectively. This trend can be attributed to the direct correlation between light intensity and the efficiency of the [2+2] cycloaddition reaction between acrylamidylpyrene - functionalized gelatin chains. Higher light intensities provide greater photon flux, enhancing the excitation of acrylamidylpyrene groups and accelerating covalent bond formation (43, 44).

Lastly, the material’s capacity for irradiation dose-dependent crosslink density was investigated through monitoring the evolution of *G*′ in 7.5% w/v Gel-Pyr under periodic irradiation (400–500 nm, 20 mW·cm^-^²) (**Figure 3f**). Upon initial light exposure (5 s), *G*′ rapidly increased; this *G*′ increase immediately stopped upon cessation of irradiation. A subsequent light exposure led to another sharp increase in *G*′, confirming the system’s capacity for on-demand network formation. This stepwise stiffening behaviour underscores the unique advantage of Gel-Pyr over conventional systems such as GelMA, where residual free-radicals can affect continued crosslinking post-irradiation ^[25]^. Such control over network formation offers a potential system for mimicking dynamic microenvironments and studying mechanotransduction, enabling applications where staged mechanical cues are essential.

These findings highlight the tunability of the Gel-Pyr system, where adjusting concentration, temperature, light intensity and irradiation duration can control gelation kinetics without the need for photoinitiators or radical-based mechanisms. This feature is particularly advantageous for 3D extrusion bioprinting which requires mild and cytocompatible crosslinking conditions. Moreover, the ability to achieve rapid crosslinking through short exposures to high (yet cytocompatible) light intensities can substantially reduce fabrication time, especially when printing multi-layered constructs, thus improving overall efficiency and throughput.

One of the important characteristics of hydrogels for 3D extrusion bioprinting is the flow behaviour, which dictates their ability to deform and flow under applied stress. Flow curves (viscosity *vs* shear rate) provide essential insights into the material’s response to shear forces(31, 45). To further investigate the rheological characteristics of the Gel-Pyr system, the viscosity was measured as a function of flow rate (0.01 s^-1^ to 1000 s^-1^) at 25°C for a range of concentrations (3.5, 5.0, 7.5% w/v, **Figure 4a**). Under nearly all conditions, viscosity decreased with increasing shear rate, corresponding to a non-Newtonian shear-thinning behaviour of Gel-Pyr. This property facilitates fluid flow as shear rate increases, advantageous for bioprinting and injection as it facilitates smooth extrusion under shear while enabling rapid structural recovery post-deposition, ensuring shape fidelity and mechanical integrity (45). However, the data for low shear rates (0.01 to 0.1 s^-1^) showed that all samples exhibited a slight increase in viscosity. This may be attributed to initial closeness and rearrangement of the Gel-Pyr chains, facilitating π-π stacking between the pyrene side-chains and interactions between the gelatin molecules. In addition, the multi-helical structures within the hydrogel can create resistance to flow when subjected to primary weak external forces. It is assumed that increasing the shear rate disrupts these physical interactions, leading to deformation of the polymeric chain segments, and a subsequent decrease in viscosity (46). The viscosity of the hydrogels increased at higher concentrations, with the highest viscosity observed for 7.5% w/v Gel-Pyr at 25°C. Shear-thinning behaviour is a desirable property for extrusion bioprinting, as the shear rate rises when the bioink passes through the nozzle (47). However, compared to pure GelMA, a study by Rastin *et al*. reported that even though 5.0% w/v GelMA shows shear-thinning behaviour, its low viscosity causes a lack of printability (48). Therefore, the hydrogel’s viscosity should neither be excessively high to impede flow, which could lead to nozzle clogging, nor too low to lack fidelity. The shear-thinning properties can be quantitatively analysed by applying a power-law regression (Ostwald-de Waele model) to the linear region of viscosity-shear profile (between 0.1 to 100 s^-1^ for 3.5% and 5.0% w/v, between 1 to 1000 s^-1^ for 7.5% w/v) (34). The equation is given as:

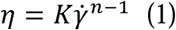

where is the viscosity, *γ̇* is the shear rate, *K* and *n* are fluid consistency index and power law index, respectively. Values of *n*<1 represents shear-thinning behaviour. The values determined through curve fitting are listed in Table 1. All tested concentrations demonstrated a pronounced shear-thinning profile, characterized by *n* values of 0.1 for Gel-Pyr 5.0% and 7.5% w/v and 0.3 for 3.5% w/v, respectively. The relatively weaker shear-thinning profile of 3.5% w/v correlates with its lower viscosity, resulting in reduced structural fidelity. The values for Gel-Pyr 7.5% w/v at range of temperature from 20 to 30 °C are listed in **Table S1 Supplementary documents**.

**Figure 4.**
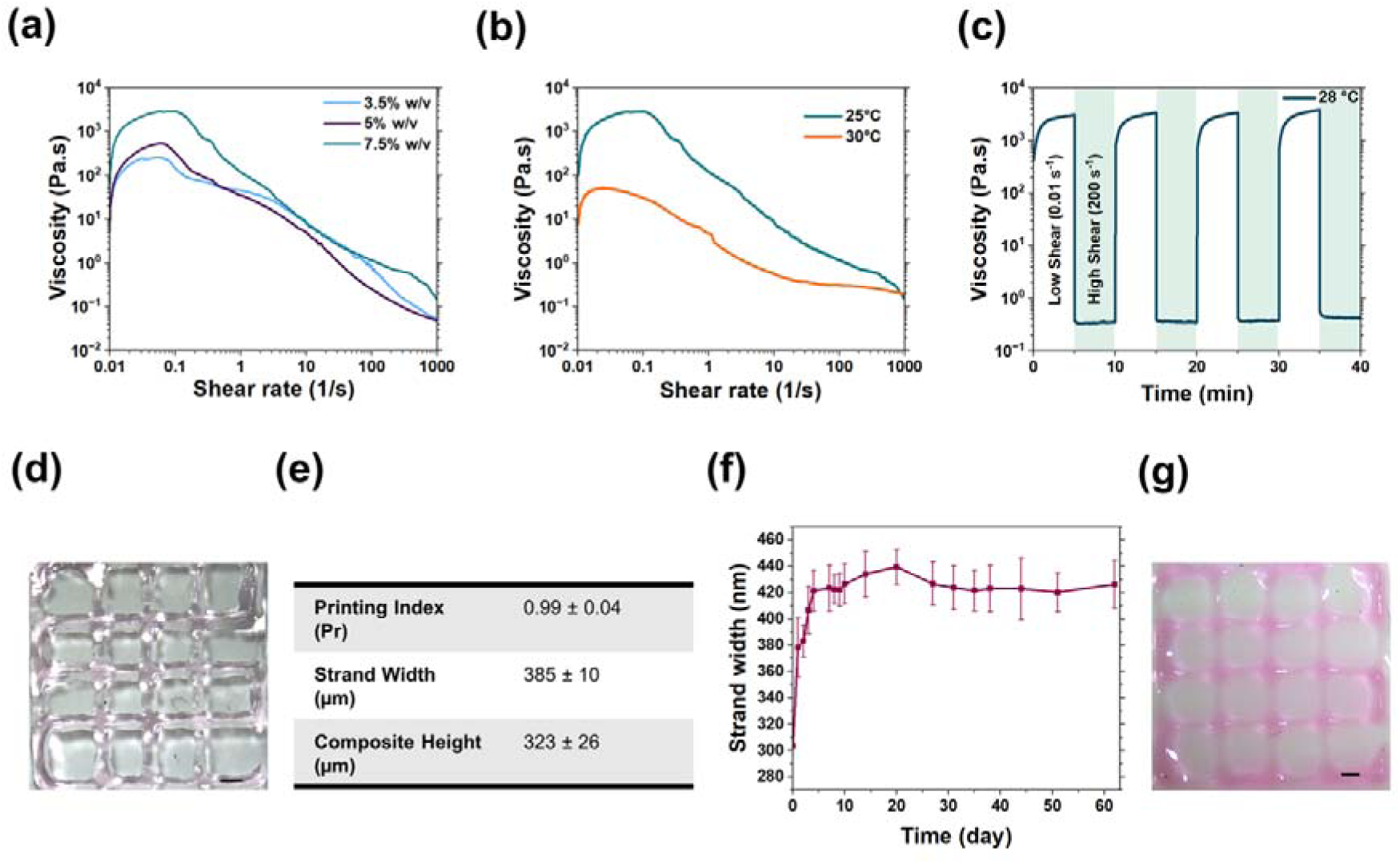
Flow behaviour and extrusion bioprinting of Gel-Pyr hydrogels. a) Viscosity under shear-rate sweep of Gel-Pyr 3.5% w/v, 5.0% w/v and 7.5% w/v at 25°C. b) Viscosity versus shear rate of Gel-Pyr 7.5% w/v at 30°C, with 5 min pre-incubation at 4°C to mimic the bioprinting condition. c) Viscosity recovery of Gel-Pyr 7.5% w/v under alternating low (0.01 s^-1^) and high (800 s^-1^) shear rate at 28°C. d) 3D bioprinted crosshatch structure fabricated via extrusion-based bioprinting using Gel-Pyr 7.5% w/v at 28°C, with a constant extrusion pressure of 40 kPa and a printing speed of 11 mm·s^-1^. Scale bar: 600 µm. (e) Semi-quantitative analysis of the bioprinted structure parameters, calculated using ImageJ (n=3, mean ± SE). (f) Swelling kinetics of bioprinted Gel-Pyr (7.5% w/v) immersed in DMEM culture medium over 62 days (n=3, mean ± S.E.M.). (g) Representative image of a swellen bioprinted structure at day 27. Scale bar = 600 µm.

**Table 1.** Power-law fitting parameters for Gel-Pyr precursor solutions. Consistency index (K), power law coefficient (n) and zero-shear rate (η_0_) for non-irradiated Gel-Pyr precursor, where K and n are derived from linear regression of viscosity-shear rate plots at 25°C.

| Concentration (% w/v) | n | K | η<sub>0</sub> (Pa·s) |
| --- | --- | --- | --- |
| 3.5% | 0.3 | 36.3 | 255 |
| 5.0% | 0.1 | 33.1 | 530 |
| 7.5% | 0.1 | 82.2 | 2893 |

According to the flow behaviour, at 3.5% w/v, the viscosity was 146 ± 24 Pa·s, corresponding to low (0.1 s^-1^) shear rates. At 5.0% w/v, the viscosity was 326 ± 150 Pa·s, and it markedly increased to 29000 ± 1200 Pa·s at 7.5% w/v. Primarily, increasing the polymer concentration led to a substantial rise in viscosity, particularly at low shear rates, highlighting its impact on rheological behaviour.

### 2.3 Gel-Pyr bioink for photoinitiator-free 3D bioprinting

The combination of tuneable stiffness, photoinitiator-free photo-crosslinking, viscoelasticity, and shear-thinning behaviour positions Gel-Pyr as a promising candidate for 3D bioprinting and tissue modelling applications. Physiologically, an ideal bioink should provide a suitable microenvironment to support essential cellular functions such as proliferation, differentiation and tissue formation, while maintaining cell integrity and survival post printing (49, 50). The primary processes include the steady extrusion of ink through the nozzle, the movement of the nozzle along predefined paths for layerwise material deposition, and final solidification. To explore this potential, a qualitative approach was employed to determine the optimal printing parameters using the 7.5% w/v Gel-Pyr formulation. Successful 3D bioprinting requires careful tuning of all parameters to ensure both good printability and high cell viability. In this study, the optimized parameters for continuous and uniform filament formation included a needle inner diameter (ID) of 150 μm, a needle length of 6.35 mm, and a printing pressure of 40 kPa. This setup specifically addresses the significant impact of dispensing pressure on cell viability (51).

To further refine the printing process, the effect of temperature on the flow behaviour of Gel-Pyr 7.5% w/v was assessed across a temperature range of 15°C to 30°C **(Figure S5, Supplementary Information)**. The theoretical shear stress and shear rates experienced by the hydrogel precursor under different conditions were determined using the Metzner–Reed (52) and power-law models (**Eq. (1), (2) and (3) Supplementary Information**). The bioprinting shear rate, viscosity, and shear stress are summarized in **Table 2**.

**Table 2.**
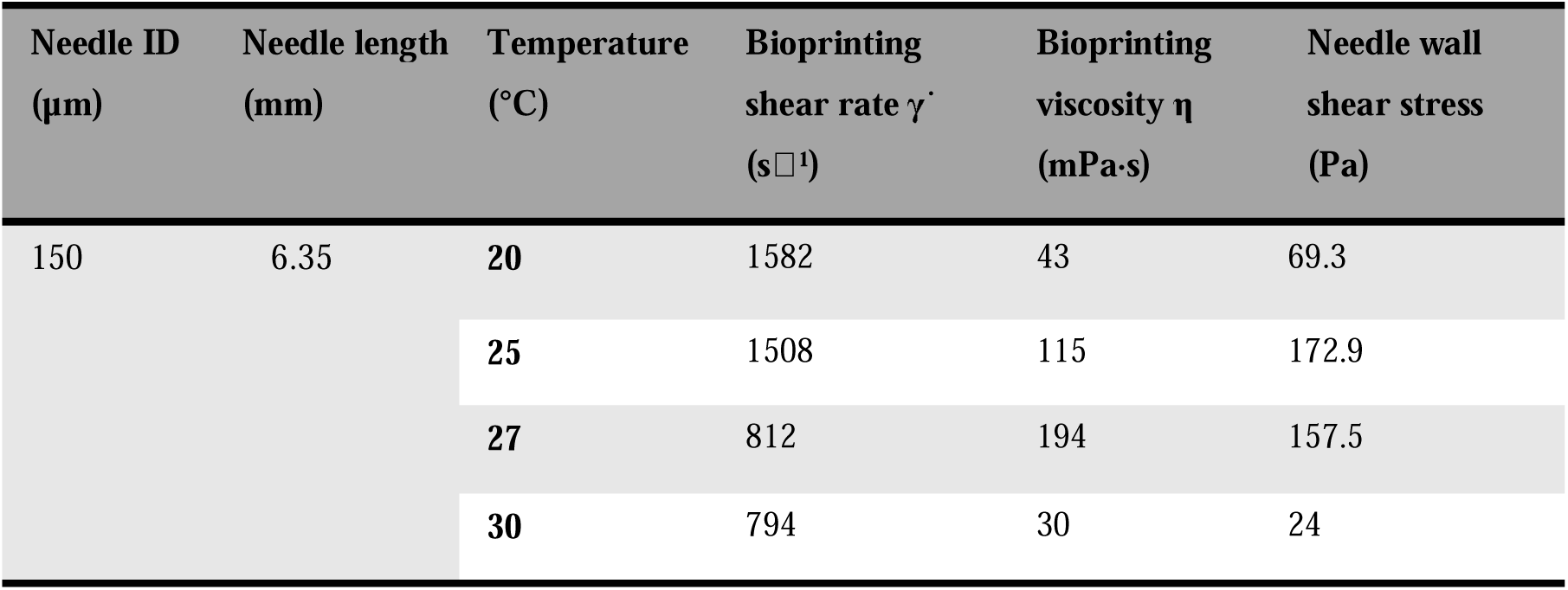
Bioprinting parameters for Gel-Pyr precursors. The bioprinting shear rate, corresponding viscosity, and shear stress for Gel-Pyr 7.5% w/v were calculated at temperatures ranging from 20°C to 30°C using the Metzner–Reed model and the power law model.

At temperatures below 27°C, frequent nozzle blockages were observed, likely due to increased viscosity which impeded extrusion. At 27°C, uniform strand extrusion was achieved, albeit requiring higher printing pressure. In contrast, at temperatures above 27°C, strand extrusion remained uniform with reduced pressure requirements. As demonstrated in **Figure 4b** and Table 2, Gel-Pyr 7.5% w/v at 30°C exhibited a pronounced shear-thinning profile (n = 0.26) and a reduction in needle wall shear stress (24 Pa), further supporting its suitability for extrusion-based bioprinting.

The shear recovery and shape retention of Gel-Pyr 7.5% w/v were examined by applying alternating shear rates (high = 800 s^-1^, low = 0.01 s^-1^) to replicate the conditions experienced during bioprinting (**Figure 4c**). The selection of the high shear rate of 800 s^−1^ was based on the averaged theoretical shear rate estimated under optimal extrusion conditions (Table 2). When the shear rate increased from 0.01 to 800 s ¹, the viscosity significantly decreased from 500 ± 230 Pa·s to 0.30 ± 0.03 Pa·s; however, upon removal of the high shear rate, the viscosity rapidly recovered. Practically, the dynamic inter-strand interactions underwent transient disruption during high shear application, facilitating extrusion, but gradually reformed through hydrogen bonding under low shear. The viscosity recovery plays a crucial role in governing the transition from fluid-like behaviour to elasticity following shear-induced deformation (53). These findings highlight that the material properties were effectively restored to their pre-deformation state, ensuring high shape retention of the printed filaments (54).

The filament fidelity was further assessed by measuring strand diameters in crosshatch scaffolds (**Figure 4d**). Within the tested printing conditions (Gel-Pyr 7.5% w/v, pressure= 40 kPa, needle ID = 150 μm, temperature = 28°C), the printed strands exhibited a mean width of 385 ± 10 μm. This width aligns with the optimal range for cell-laden constructs, as previous studies have identified 200–400 μm as the ideal strand width for enhancing nutrient/waste exchange and facilitating cell migration (55). Additionally, printability was quantitatively assessed based on pore morphology using the printability index (*Pr*), calculated according to Eq. (2). The *Pr* provides a numerical measure of the squareness of interconnected channels in the crosshatch scaffold and is defined as:

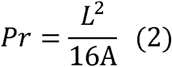

where *L* represents the pore perimeter and *A* denotes the pore area. A *Pr* value of 1 corresponds to an ideal square pore shape, while *Pr* < 1 and *Pr* > 1 indicate round and irregular pore geometries, respectively (20, 34). The optimized Gel-Pyr precursor demonstrated a *Pr* value of 0.99 ± 0.04 (**Figure 4e**), indicating near-perfect square pore morphology, which correlates with complete and rapid viscosity recovery kinetics.

Subsequently, the stability of the bioprinted constructs was assessed during 62 days of incubation in DMEM under physiological conditions (**Figure 4f**). Following an initial swelling phase, during which the average strand width increased from 303 μm at day 0 to 421 μm at day 4, no further significant changes in strand dimensions were observed. Importantly, the overall construct architecture remained intact throughout the study period, minimizing post-fabrication deformation (**Figure 4g**). These results demonstrate the excellent long-term structural stability of the constructs and highlight their potential for prolonged tissue culture applications.

### 2.4 Gel-Pyr is a biocompatible hydrogel, with cells well-distributed within a 3D environment

The biocompatibility of hydrogels is a critical parameter determining their suitability for applications in tissue engineering, regenerative medicine and advanced *in vitro* biological models(55). To evaluate the cytocompatibility of Gel-Pyr, human bone marrow-derived mesenchymal stromal cells (BM-MSCs) were directly encapsulated within 7.5% w/v Gel-Pyr hydrogels, photocrosslinked under visible light for 3 min, and cultured for 7 days. Confocal microscopy confirmed efficient cell encapsulation and homogeneous cell distribution throughout the hydrogel volume **(Figure 5a-d)**. Encapsulated BM-MSCs adopted a well-spread, elongated morphology, suggesting that the Gel-Pyr matrix provides a permissive microenvironment that supports cell adhesion and cytoskeletal organization. 3D reconstructions and depth-coded projections further confirmed homogeneous cellular organization throughout the constructs, demonstrating that the rapid gelation kinetics of Gel- Pyr preserve spatial cell distribution while enabling the fabrication of structurally uniform, cell-laden hydrogels.

**Figure 5.**
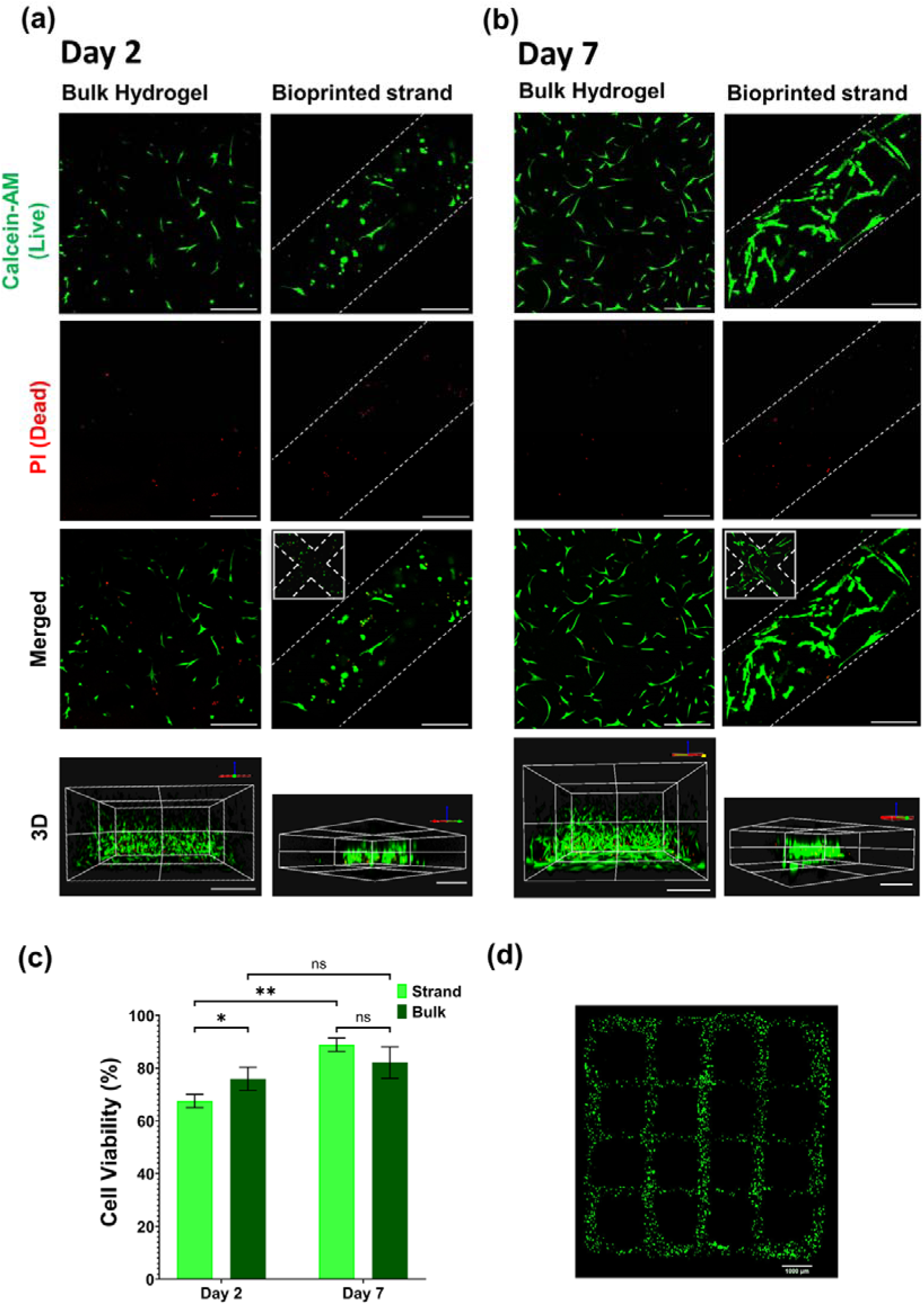
Cell viability of BM-MSCs encapsulated in bulk and bioprinted Gel-Pyr hydrogels. Live/dead staining was performed using calcein AM (green, live cells) and propidium iodide (red, dead cells). (a,b) Representative confocal micrographs of encapsulated BM-MSCs cultured for 2 and 7 days, respectively. c) Quantitative analysis of cell viability in bulk and bioprinted Gel-Pyr hydrogels over the culture period. Data are presented as mean ± S.E.M. (n = 3); *P < 0.05, **P < 0.01 (two-way ANOVA). d) Representative confocal image showing the distribution of viable BM-MSCs throughout a bioprinted crosshatch construct. Scale bars: 300 μm in (a,b) and 1000 μm in (d).

Viability of BM-MSCs in Gel-Pyr was then assessed via live/dead staining 2 and 7 days post-encapsulation. Cell survival was studied under the following conditions: (i) cells encapsulated in bulk gel and (ii) cells embedded in 3D bioprinted crosshatch constructs.

Images at day 2 showed that the BM-MSCs had begun to elongate, with more pronounced morphological changes observed by day 7, where cells displayed spreading, and more defined spindle-like shape, suggesting appropriate adaptation and favourable interaction with the hydrogel (**Figure 5a and b**). Quantitative analysis revealed that cell viability within the bulk scaffold was 76 ± 4 and 82 ± 6% at days 2 and 7, respectively, as indicated by the predominant green calcein fluorescence (**Figure 5c**). The bioprinted constructs resulted in 68 ± 2% and 89 ± 3% cell viability at the same time points. The statistically significant reduction in viability when comparing printed structures to bulk hydrogels at day 2 can be attributed to the shear stress experienced by cells during extrusion-based bioprinting, which has been shown to impact membrane integrity and induce apoptosis (56). However, by day 7, viability in the bioprinted constructs increased significantly, reaching levels comparable to those in bulk hydrogels suggesting that the initially stressed cells were able to recover over time, presumably due to the supportive microenvironment provided by the hydrogel. Notably, the bulk system exhibited no significant changes in cell viability between days 2 and 7.

Overall, the results suggest that Gel-Pyr provides a favourable 3D microenvironment that supports cell survival. Furthermore, the enhanced cell viability at day 7 in bioprinted constructs indicates not only the hydrogel’s cytocompatibility but also its ability to promote cell proliferation, likely due to improved nutrient and oxygen diffusion within the fine bioprinted strands.

Free radicals generated during photoinitiator-mediated crosslinking are known to induce intracellular oxidative stress and subsequent DNA damage, and changes in key signalling pathways in encapsulated cells (57, 58). To evaluate whether the photoinitiator-free gelation mechanism of Gel-Pyr could alleviate these effects, we compared Gel-Pyr with GelMA as the widely adopted photocrosslinking condition reported for tissue engineering applications (59–61). To ensure a comparable evaluation, the mechanical properties of GelMA were first characterized and adjusted to closely match those of Gel-Pyr (**Figure S6- Supplementary Information**). As shown in **Figure 6c,d**, BM-MSCs encapsulated in GelMA (6% w/v + 0.5% w/v LAP) exhibited substantially elevated CellROX fluorescence intensity (38.9 ± 2.8), indicating increased ROS accumulation during photocrosslinking. In contrast, Gel-Pyr (7.5% w/v) showed significantly reduced ROS levels (7.8 ± 1.6), suggesting that the absence of photoinitiator effectively minimized oxidative stress during gelation. While excessive ROS can induce oxidative DNA damage, mitochondrial dysfunction and apoptosis, sublethal ROS levels are also sufficient to impair key MSC functions despite the cells remaining viable. For example, ROS accumulation reduces MSC stemness and self-renewal capacity, limiting their regenerative potential (62). It also promotes premature senescence, with senescent MSCs establishing a feed-forward loop that sustains ROS-associated stress and amplifies its downstream effects (63–65). Elevated ROS has also been shown to skew multi-lineage differentiation, favouring adipogenic over osteogenic lineages (66, 67). Beyond these intrinsic effects, oxidative stress compromises MSC homing and engraftment and alters their paracrine and immunomodulatory functions (68, 69). Collectively these examples highlight the potential detrimental effects of ROS exposure, specifically on the very processes by which the MSCs are expected to support tissue repair and regeneration. Given the relatively robust antioxidant defences and stress tolerance of MSCs, the benefits of reducing ROS exposure observed here are likely to be at least as relevant, and potentially more significant, for other regenerative medicine cell types that are more susceptible to oxidative stress (70).

**Figure 6.**
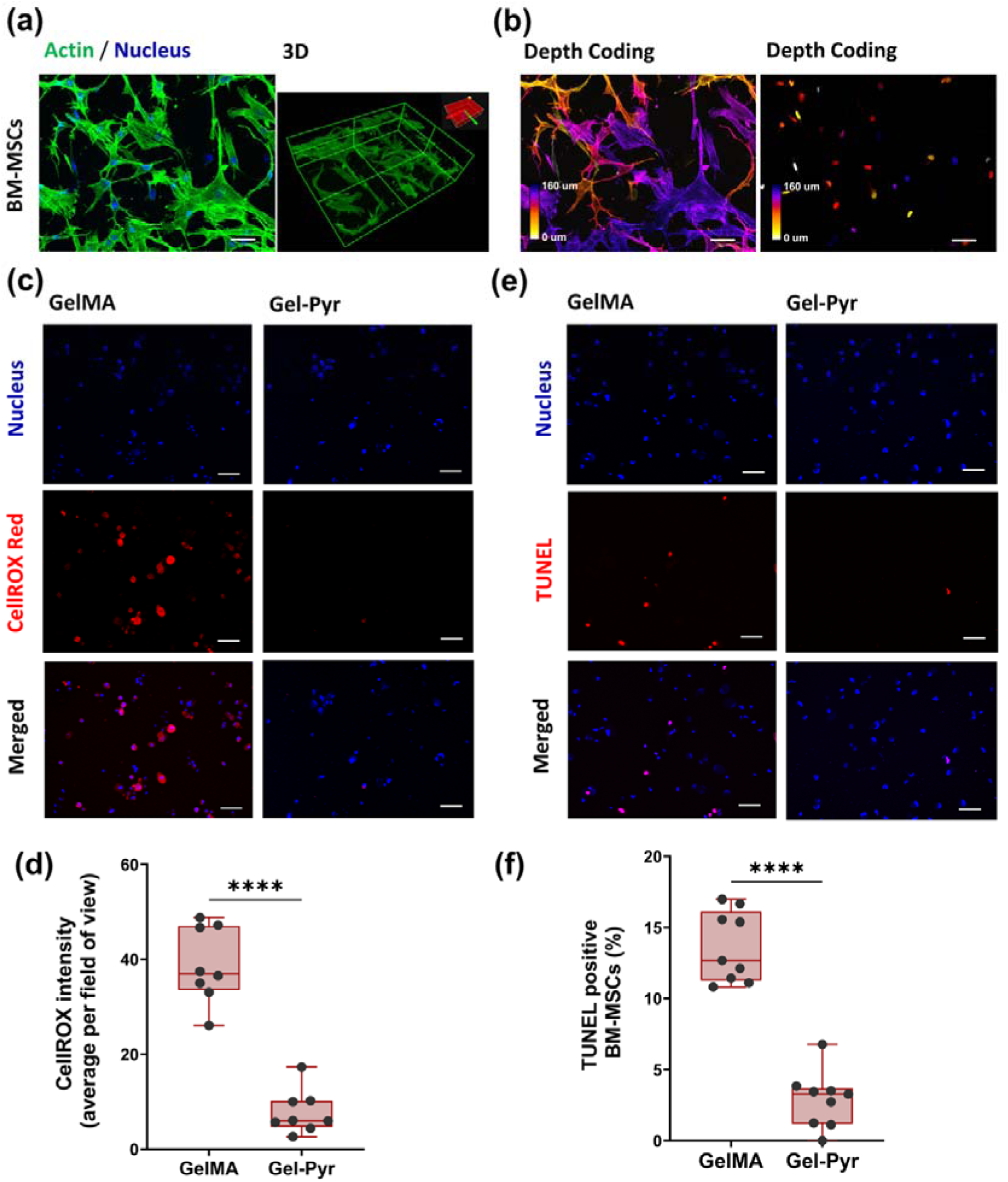
Cellular compatibility and cytoprotective effects of Gel-Pyr hydrogels. (a) Immunofluorescence images and 3D reconstruction of BM-MSCs encapsulated within 7.5% w/v Gel-Pyr for 7 days. F-actin is shown in green and nuclei in blue. (b) Depth-coded projections showing the spatial distribution of the BM-MSC cytoskeleton (left) and nuclei (right) throughout the hydrogel construct. (c) Representative CellROX™ Red staining images of BM-MSCs encapsulated in GelMA and Gel-Pyr hydrogels, indicating intracellular reactive oxygen species (ROS) generation 2 min after blue-light exposure. (d) Quantification of CellROX fluorescence intensity in BM-MSCs encapsulated in GelMA and Gel-Pyr hydrogels, demonstrating significantly reduced ROS levels in Gel-Pyr. (e) Representative TUNEL staining images of BM-MSCs encapsulated in GelMA and Gel-Pyr hydrogels after 48 h of culture. (f) Quantification of TUNEL-positive BM-MSCs, demonstrating significantly reduced apoptosis in Gel-Pyr compared with GelMA after 48 h. Data are presented as mean ± S.E.M. (n = 9); ****P < 0.0001 (unpaired two-tailed Student’s t-test). Scale bars: 50 μm.

Notably, the ROS profiles closely corresponded with the TUNEL results, where GelMA- encapsulated BM-MSCs displayed markedly higher DNA fragmentation, with 13.6 ± 0.8% TUNEL-positive cells compared with 2.9 ± 0.6% in the Gel-Pyr group (**Figure 6e, f**). This strong correlation between CellROX and TUNEL analyses supports radical-induced oxidative stress as a key driver of the genomic damage observed in GelMA-encapsulated cells. Conversely, the lower ROS levels in Gel-Pyr were accompanied by minimal TUNEL positivity, demonstrating improved preservation of genomic integrity following encapsulation. Collectively, these results highlight the advantage of the photoinitiator-free Gel-Pyr system in mitigating oxidative and genotoxic stress, thereby providing a more cytocompatible microenvironment for BM-MSCs encapsulation than conventional photocrosslinked hydrogels.

## 3 Conclusion

In this study we developed a catalyst-free photo-crosslinkable hydrogel system based on gelatin functionalized with acrylamidylpyrene chromophore, termed Gel-Pyr. This system presents a significant advancement over previous hydrogel platforms by eliminating the need for added photoinitiators, which are commonly associated with radical-mediated crosslinking, cytotoxicity, and additional formulation complexity. Harnessing the capability of acrylamidylpyrene to undergo [2+2] cycloaddition, rapid polymerization was achieved under visible light (400–500 nm) irradiation, facilitating a high degree of crosslinking in under 50 s at moderate irradiation intensities without generating reactive radicals.

Gel-Pyr exhibited suitable viscoelastic and mechanical properties, with tuneable pre-gelation characteristics and extended stability for over 32 days, making it highly suitable for both 2D and 3D bioengineering applications. In addition, its ability to undergo on-demand network formation through light irradiation offers precise spatiotemporal control over crosslinking, further expanding its versatility in complex tissue fabrication. Notably, Gel-Pyr demonstrated shear-thinning behaviour, establishing its utility as a robust bioink for extrusion-based 3D bioprinting. Its optimized flow characteristics facilitated the fabrication of crosshatch structures with fine strands (<400 µm diameter), enabling enhanced nutrient and gas exchange to support encapsulated cell viability and function within a 3D architecture. Moreover, Gel-Pyr provided a highly biocompatible microenvironment with significantly reducing intracellular ROS generation and DNA damage compared with GelMA. This cytoprotective effect was accompanied by high cell viability (>80%) and sustained cellular proliferation within bioprinted constructs.

Overall, this initiator-free and low-radical generating strategy, offering rapid spatiotemporal control of crosslinking, streamlines the system and substantially enhances cytocompatibility. As a result, it represents a key innovation that addresses major limitations of conventional photocurable platforms and serves as a more versatile alternative to widely used GelMA, particularly in 3D bioprinting applications.

## 4 Experimental Section

### Materials

Gelatin (type A, bloom strength: 300 g), pyrene-1-aldehyde, malonic acid, pyridine, piperidine, dimethylformamide (DMF), 1-ethyl-3-(3-dimethylaminopropyl)carbodiimide hydrochloride (EDC·HCl), *N*-hydroxysuccinimide, dichloromethane, penicillin streptomycin (P/S), Bovine serum albumin (BSA) (A7906), Triton X-100 (X100), and Tween 20 (P1379) were purchased from Sigma-Aldrich. Advanced Dulbecco’s modified Eagle medium (DMEM) (10567–014), Neurobasal-A medium (10888022), L-Glutamine (25030081), B-27 (17504044), TrypLE™ Express (12605-028), Calcein AM dye (C1430), propidium iodide dye (P1304MP), CellROX Oxidative Stress Reagents (C10443), Click-iT™ Plus TUNEL Assay Kit (C10618), and DRAQ5™ Fluorescent Probe Solution (5 mM) (62251) were purchased from Thermo Fisher Scientific. were purchased from Sigma-Aldrich. ActinRed dye was purchased from Life Technologies (R37112). Fetal bovine serum (FBS) (SFBS) was purchased from BovoGen. Dimethyl sulfoxide (DMSO) was purchased from Hello Bio (HB3262).

### Synthesis of Gelatin-Pyrene (Gel-Pyr)

Gel-Pyr macromers were synthesised through a three-step synthetic protocol. First, 1- pyrenecarboxaldehyde (5 g, 21.7 mmol) was mixed with malonic acid (4.16 g, 40.0 mmol), pyridine (26 mL), and piperidine (8 mL). This mixture was heated at 90°C for 16 hours, then poured into 200 mL of ice-cold 1 M HCl. The resulting solid was collected by filtration and dried under vacuum to afford yellow powder, 3-(1-pyrenyl)acrylic acid (yield: 5.03 g, 85.0%), and used directly in the next step. Next, 3-(1-pyrenyl)acrylic acid (1 g, 3.67 mmol) was mixed with EDC·HCl (0.84 g, 4.40 mmol) and N-hydroxy succinimide (0.51 g, 4.40 mmol) in anhydrous THF (20 mL) and anhydrous DMF (5 mL). The mixture was stirred at room temperature for 6 h then added to iced cooled distilled water (100 mL). The precipitate was filtered then dried in vacuum at 40 °C, and further purified via column chromatography using dichloromethane as the eluent and collected as a yellow powder, acryl pyrene with N-hydroxy succinimide ester (yield: 1.14 g, 84.0%). Finally, to prepare hydrogel precursors, a gelatin solution was obtained by dissolving 2 g of porcine skin gelatin type-A in distilled water (15 mL) and DMF (15 mL). Then, the obtained acryl pyrene with N-hydroxy succinimide ester (73.9 mg, 0.2 mmol) was first dissolved in DMF (2 mL) and added to the gelatin solution. The reaction was stirred at 40 °C for 24 h. The polymer was then precipitated by adding the crude mixture into acetone (300 mL). The collected polymer was subsequently dialyzed (molecular weight cut-off 1 kDa) against DI water at 40 °C for 2 days. The purified solution was freeze-dried for 3 days, yielding a pale yellow solid (yield: 1.8 g, 86.8%).

### Gel-Pyr Hydrogel Preparation

Gel-Pyr solutions were prepared at varied concentrations by weighing an appropriate amount of Gel-Pyr and dissolving it in pre-warmed PBS or low glucose DMEM. The Gel-Pyr solution was incubated in a water bath at 45-50°C until complete dissolution and disappearance of Gel-Pyr solid.

### Characterisation of Gel-Pyr hydrogel

#### UV-Vis spectrometry

The conjugation of acrylamidylpyrene to gelatin was confirmed by UV-Vis spectrometry. UV-Vis absorption spectra were acquired using Cary 4000 UV-Vis Spectrometer (Agilent Technologies, Santa Clara, CA, USA). Spectra were collected from Gel-Pyr and porcine skin gelatin as a control. To that end, both solutions were prepared at 0.05% w/v in DMSO. Using a quartz cuvette with a path length of 1.0 cm, absorbance was recorded by running a spectral scan from 300-500 nm at room temperature. Baseline correction was performed using DMSO as blank solution. Each sample was scanned three times to ensure reproducibility, and the average absorbance values were recorded.

To determine the degree of substitution, an acrylamidylpyrene -amide standard curve was employed with standard sample solutions prepared at 0, 5.43×10^-6^, 1×10^-5^, 2.1×10^-5^, 4.3×10^-5^, and 8.7×10^-5^ mol/L in DMSO. A solution of Gel-Pyr hydrogel with 0.0625% w/v concentration was also prepared in DMSO and the absorbance of all samples was recorded at 399 nm. Finally, the concentration of pyrene in the Gel-Pyr solution was calculated using linear regression analysis of the standard curve.

### Rheological characterisation

The mechanical properties and viscoelastic parameters of the Gel-Pyr precursors were investigated using a stress-controlled rheometer (MCR Physica 501, Anton Paar) equipped with a temperature-controlled lower plate and a photo-curing stage. All measurements were collected in triplicate using parallel plates (12 mm diameter, 200 μm gap) for oscillatory experiments, and 1° cone-plate (25 mm diameter, 48 μm gap) for rotational experiments. An Omnicure light source (S2000 spot UV cure system, Excelitas Technologies), emitting light in the 400-500 nm range was employed to photo-crosslink the hydrogel precursors. To prevent dehydration, the edges of each sample was sealed with paraffin oil. Experiments were conducted with Gel-Pyr samples at 3.5, 5.0, 7.5, and 10.0 % w/v in low glucose DMEM, and at temperatures ranging from 15 °C to 30 °C.

For the thermoresponsive behaviour investigations, 7.5% w/v Gel-Pyr precursors were loaded between the parallel plates and shear modulus were recorded through a series of sequential heating (4°C to 40°C) and cooling (40°C to 4°C) cycles at a ±1 °C min^-1^. The frequency sweep was performed on Gel-Pyr precursors over a frequency range of 0.1 Hz to 10 Hz at a strain of 1%. The amplitude sweep experiments were subsequently conducted at a constant frequency of 1 Hz over the strain range of 0–100% to determine the linear viscoelastic region. The rotational experiments employed to determine the shear viscosity as a function of the shear rate, where the shear rate varies from 0.01 s^−1^ to 1000 s^−1^. To investigate the self-recovery and shape fidelity a cyclic shear rate - time sweep test was performed at alternating low (0.01 s^-1^) and high (800 s^-1^) shear rates in 5-min intervals.

Crosslinking was monitored under 400-500 nm light illumination at intensities adjusted to 5, 10 and 20 mW cm^-2^ (using the radiometer RM12, Gigahertz-Optik, Germany). The samples were loaded between parallel plates at a 0.2 mm gap and conditioned at varying temperatures for 30 s followed by light exposure of 8 min to initiate the gelation and determine the photopolymerization time.

To evaluate the temporal control of visible light-induced crosslinking, 7.5% w/v Gel-Pyr precursors (prepared in DMEM) were loaded onto a rheometer stage preheated to 25°C and subjected to alternating light on/off cycles following an initial 20-second delay. Visible light (400–500 nm) at an intensity of 20 mW·cm^-^² was applied in 5 s pulses separated by 20 s dark intervals until t = 2.25 min. From t = 2.25 to 3.25 min, the light exposure was increased to 10 s intervals, followed by continuous irradiation from t = 3.25 min onward.

### Mechanical characterization

Uniaxial compression testings of the Gel-Pyr samples were conducted on rheometer under constant speed of 0.3 mm min^-1^. Gel-Pyr precursors (3.5%, 5.0% and 7.5% w/v) were dissolved in DMEM and hydrogel disks were prepared in cylindrical Teflon molds (10 mm diameter, 2 mm high) followed by crosslinking with 405 nm light irradiation (20 mW cm^-2^) for 5 min. Subsequently, samples were incubated overnight in DMEM to reach equilibrium swelling prior to testing. Briefly, for the compression testing, the hydrogel cylinders were positioned atop the stationary bottom plate, with the temperature maintained at 37 °C. To adjust the initial position of the upper parallel plate (12 mm diameter), the plate was gradually lowered to contact the gel’s surface, ensuring an applied normal force of 0.01 N. To construct the stress-strain curve, the strain was calculated using Eq. (3), where *h*_0_ is the initial height, and Δ*h* is the displacement of the top plate, while the stress was determined from the force, all recorded by instrument during compression. The compression modulus was determined from the stress-strain curve (**Figure S4- Supplementary Information**) over 10- 20% strain, corresponding to the slope in the linear region.

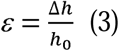

### Mass swelling and mass loss analysis

Gel-Pyr hydrogels (7.5% w/v, n = 3) were prepared either as bulk samples (10 μL droplets) or as bioprinted constructs using the optimized printing parameters. Samples were photocrosslinked by visible light irradiation (400–500 nm, 20 mW cm^-^²) for 3 min. Bulk samples were weighed immediately after gelation, while strand widths were measured in bioprinted constructs. Hydrogels were subsequently immersed in low-glucose DMEM and incubated under physiological conditions (37 °C, 5% CO, humidified atmosphere), with medium replacement every 48 h. At predetermined time points, excess medium was removed and the swollen weight (W_t_) and strand widths were recorded. For mass loss analysis, bulk samples were freeze-dried at each time point and the dry weight (d_t_) was measured. Hydrogels were monitored for up to 7 days. Apparent swelling and mass loss were calculated according to Equations (4) and (5), respectively:

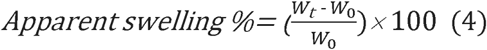

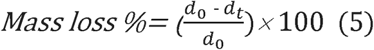

where *W*_0_ and *W_t_* represent the initial and swollen hydrogel weights, respectively, and *d*_0_ and *d_t_* correspond to the initial dry weight before swelling and the dry weight at time *t*, respectively.

### Gel fraction evaluation

Gel-Pyr hydrogels (7.5% w/v in deionized water, n = 3) were prepared by casting the precursor solution into Teflon molds (10 mm diameter, 2 mm height) and crosslinked via exposure to 405 nm light at an intensity of 20 mW cm^-^² for 5 minutes. The initial dry weight was determined (*W*_0_) and the samples were incubated in deionized H_2_O at 37°C for 48 h to allow leaching of the sol fraction, with the water refreshed after 24 h. Following equilibrium swelling, the hydrogels were freeze-dried for 24 h, and their final dry mass (*W_t_*) was measured. The gel fraction (%) was calculated using Eq. (6):

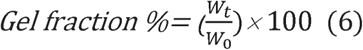

### 3D bioprinting optimisation

Gel-Pyr precursors were dissolved in prewarmed DMEM culture medium and incubated at 45-50°C to fully dissolve. The bioink was loaded into a 3 ml syringe barrel and centrifuged at 300 g for 2 min to remove any entrapped air bubbles. The mixture was then cooled on ice for 5 min before being loaded onto the BioScaffolder 3.2 (GeSIM). To maintain a consistent temperature throughout the printing process, a custom Peltier jacket was used to enclose the syringe. The bioink was allowed to equilibrate at the desired temperature for ∼40 min before bioprinting. The 30 G straight stainless-steel printing nozzle (150 µm inner diameters, 6.35 mm length) (Nordson, Australia) was attached to the syringe. The bioink was then printed onto 12 mm glass coverslips under an extrusion pressure of 40 kPa and at a printing speed of 11 mm/s. The scaffolds were printed in a crosshatch pattern, consisting of three and five layers, with final dimensions of 9 mm × 9 mm. Following the deposition of all layers, the scaffolds were irradiated with visible light (400-500 nm), at 20 mW cm^-2^ intensity for 3 min. Micrographs of the printed constructs were taken with stereo zoom microscope (SMZ800N, Nikon).

### Mathematical model

The shear stress and shear rate generated by non-Newtonian fluid passing through the nozzle during 3D bioprinting were modelled along with Metzner–Reed and Ostwald-de Waele models, as detailed in the SI.

### Gel-Pyr hydrogel cell biocompatibility

#### Cell Culture

Human bone marrow-derived mesenchymal stem cells (BM-MSCs) were obtained from RosterBio (Frederick, MD, USA). BM-MSCs were cultured in growth medium consisting of low glucose DMEM, supplemented with 10% v/v FBS, 100 U mL^-1^ penicillin, and 100 µg mL^-1^ streptomycin (1% pen/strep). Cells were maintained in a humidified incubator at 37 °C and 5% CO, passaged upon reaching ≈80% confluency, and used up to passage 6.

### 3D Cell Culture in Bulk Gel-Pyr hydrogels

To synthesise cell-laden hydrogels, BM-MSCs were trypsinised and resuspended in a 7.5% w/v Gel-Pyr precursor solution in DMEM at a density of 1.5×10^6^ cells/ml. A 10 μl droplet of hydrogel-cell mixture was loaded on a glass bottom dish and irradiated with visible light (λ= 400-500 nm, I=20 mW cm^-2^) for 3 min to induce gelation. Bulk cell-laden hydrogel constructs were immersed in 2 ml low glucose DMEM supplemented with 10% FBS and 1% pen/strep and incubated for 2 and 7 days at 37°C with 5% CO_2_. The medium was refreshed every other day.

### 3D bioprinting cell-laden Gel-Pyr hydrogel

For 3D extrusion-based bioprinting, the bioink including 1.5×10^6^ cells/ml hydrogel was transferred to a pre-sterilized syringe and incubated at room temperature for 5 min. Then the syringe was installed on the bioprinter and incubated at 29°C for 40 min to equilibrate. Afterwards, the printing process built a five layered crosshatch structure with 9 mm diameter and 1 mm thickness on a sterilized 12 mm glass coverslip (using 30 G needle diameter and 65 kPa extrusion air pressure, with the bioprinting speed of 11 mm/s). The 3D bioprinted constructs were immediately crosslinked via 400-500 nm light irradiation for 3 min. Finally, the constructs were placed in 6-well culture plates containing 4 ml growth medium and incubated at 37°C and 5% CO_2_ atmosphere. Three separate constructs per time point were used in this experiment.

### Live/dead cell assay

BM- MSCs viability within either bulk or 3D bioprinted hydrogel constructs was assessed via Live/Dead assay. The Calcein AM (2 µM in PBS) and propidium iodide (PI) (4 µM in PBS) reagents were prepared as per manufacturer’s protocol. Briefly, cell-laden hydrogels were washed with PBS for 15 min and incubated with 400 μl staining solution for 20 min at 37°C. The samples were rinsed thrice with PBS and imaged using a confocal microscope (Olympus FluoView FV3000, Australia). All resultant z-stack images used a step size of 10 µm at a specific depth corresponding to the construct thickness. Images were analysed using Fiji (ImageJ, 1.52d) software. Viability was quantified as the ratio of live cells to total cells counted, using three representatives at each time point.

### Immunofluorescence staining

BM-MSCs encapsulated within Gel-Pyr hydrogels were fixed with 4% w/v paraformaldehyde in PBS for 15 min at day 7 of culture. Samples were permeabilized with 0.3% Triton X-100 for 10 min and subsequently blocked with 1% w/v BSA for 1 h. Actin was stained with ActinRed reagent (Life Technologyes, 1 drop mL^-1^), and the nucleus was stained with DRAQ5 (Thermo Fisher Scientific). Following PBS washing, fluorescence images were acquired using a confocal laser scanning microscope (Olympus FluoView FV3000, Australia). Resultant z-stack images used a step size of 1 µm at a specific depth corresponding to the construct thickness. Images were analysed using Fiji (ImageJ, 1.52d) software.

### Intracellular ROS measurement

Photoinduced intracellular ROS generation was quantified using CellROX™ Orange Reagent (Thermo Fisher Scientific). BM-MSC-laden Gel-Pyr (7.5% w/v) and GelMA (6% w/v, 0.5% w/v LAP) hydrogels were photocrosslinked for 3 min under visible light irradiation (400–500 nm, 20 mW cm^-^²). Two minutes after irradiation, samples were incubated with 5 μM CellROX™ Orange Reagent for 30 min at 37 °C in the dark and subsequently counterstained with DRAQ5 for 20 min. Following PBS washing, fluorescence images were acquired using a confocal laser scanning microscope (Olympus FluoView FV3000, Australia) and analysed to determine intracellular ROS levels.

### Cell apoptosis measurement

The click-iT TUNEL Alexa Flour Imaging Assay Kit (Thermo Fisher Scientific) was used to detect DNA fragmentation in BM-MSCs encapsulated in Gel- Pyr (7.5% w/v) and GelMA (6% w/v, 0.5% w/v LAP) hydrogels after 48 h, following the manufacturer’s protocol. Samples were counterstained with DRAQ5 to label cell nuclei. Each step was followed by thorough PBS washing and the samples were observed using a confocal laser scanning microscope (Olympus FluoView FV3000, Australia). Apoptosis was quantified as the percentage of TUNEL-positive cells relative to the total number of nuclei.

## Supporting Information

Supporting Information is available from the Wiley Online Library

## Funding

Australian Research Council Future Fellowship Scheme (No. FT200100880 to TFN & No. FT240100146 to JEF), A*STAR in the form of the Career Development Fund (No. 23-8313GU to XYO), the Australian Research Council Discovery Project Scheme (No. DP250104378 to HCP & JSF).

## Author contribution

Conceptualization: J.S.F, T.F.S, V.X.T., S.E.

Methodology: S.E, J.S.F, T.F.S, J.E.F,

Investigation: S.E., D.Z., X.Y.O

Visualization: S.E.

Supervision: J.S.F., T.F.S., V.X.T.

Writing-original draft: S.E.

Writing: review & editing: J.S.F., T.F.S., J.E.F., V.X.T., H.C.P., S.E.

## Conflict of Interest

The authors declare no conflicts of interest

## Data Availability Statement

Data for this study will be made available from the corresponding authors upon reasonable request.

## Supplementary Materials

### Determination of Pyrene Concentration in Gel-Pyr

A standard curve was established using pyrene-amide dissolved in DMSO at concentrations of 0, 5.43 × 10, 1 × 10, 2.1 × 10, 4.3 × 10, and 8.7 × 10 mol/L. The absorbance spectra of these solutions were recorded using a UV-Vis spectrophotometer (Agilent Technologies, Santa Clara, CA, USA) in a 1.0 cm path length quartz cuvette, scanning from 300 to 500 nm at room temperature. Baseline correction was applied using DMSO as the reference. The absorbance at 399 nm was used to generate a calibration curve (Figure S1).

To determine the pyrene concentration in Gel-Pyr, a hydrogel sample at 0.0625% w/v was prepared, and its absorbance was recorded at 399 nm under identical conditions. The concentration of pyrene in the hydrogel was then quantified using the linear regression equation derived from the standard curve. All measurements were performed in triplicate to ensure accuracy and reproducibility.

**Fig. S1.**
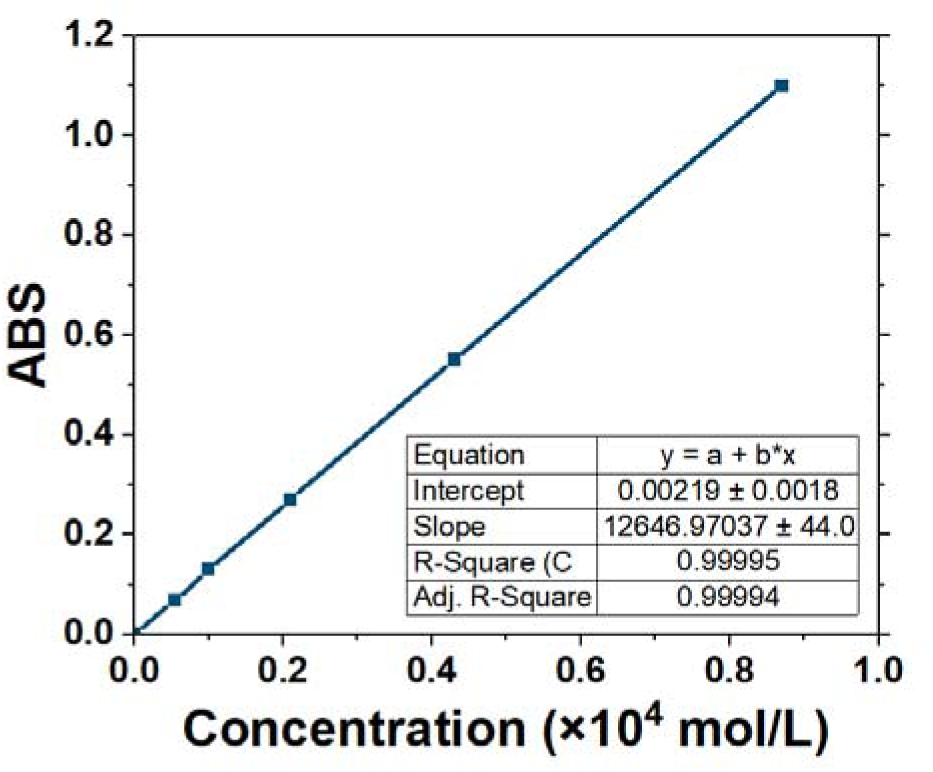
Calibration curve to determine the degree of substitution of pyrene in Gel-Pyr (from UV-Vis absorbance of the small molecule pyrene-amide).

### Rheological Characterisation

#### Physical gelation as a function of time

Due to its temperature sensitivity and time-dependent behaviour, the physical gelation process of Gel-Pyr was investigated by maintaining a 5.0% w/v solution at a constant temperature of 20 °C for 90 minutes (Figure S2). The storage modulus (G’) exhibited a significant increase within the first 15 minutes, reaching approximately 600 Pa. Following this rapid initial gelation, G’ continued to increase gradually before stabilizing after 90 minutes, indicating the completion of the gelation process.

**Fig. S2.**
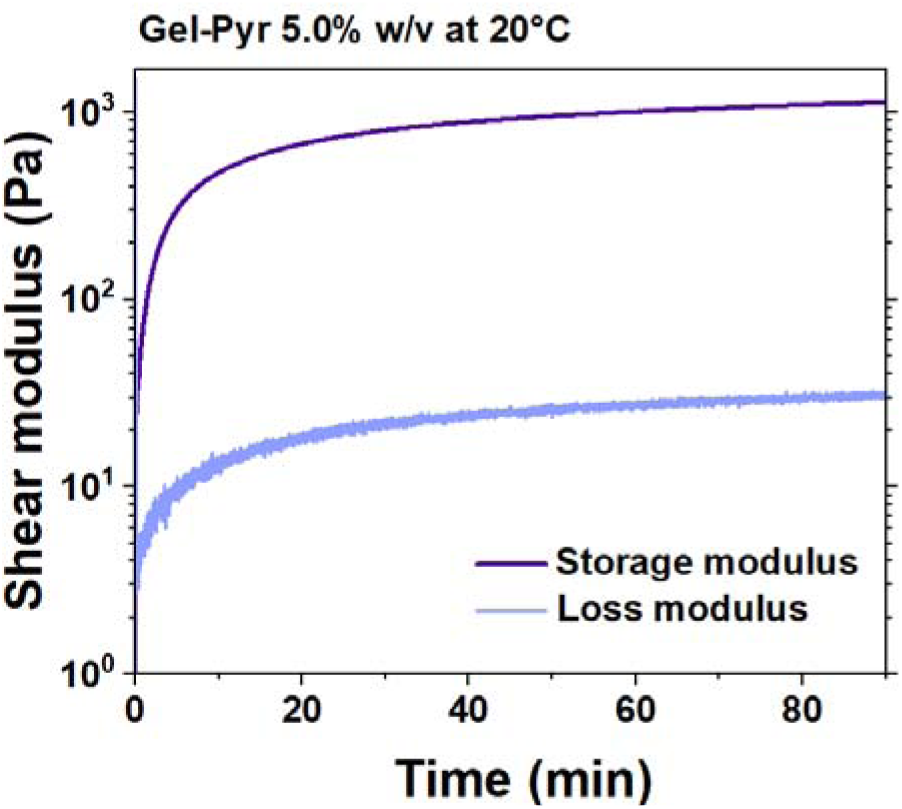
Physical gelation as a function of time at 20 °C.

### Photo-rheology of Gel-Pyr at different concentrations

To investigate the effect of concentration and temperature on the shear modulus, in-situ photo-rheology was performed for 8 min using Gel-Pyr at concentrations of 3.5%, 5.0%, and 10.0% (w/v) across a temperature range of 15–27°C. Samples were equilibrated for 30 s, and subsequently irradiated with blue light (λ = 400–500 nm, I = 20 mW·cm^-^²). As shown in Figure S3, an increase in hydrogel concentration and a decrease in temperature resulted in a higher storage modulus. These findings suggest that the mechanical properties of Gel-Pyr can be tailored based on specific application requirements.

**Fig. S3.**
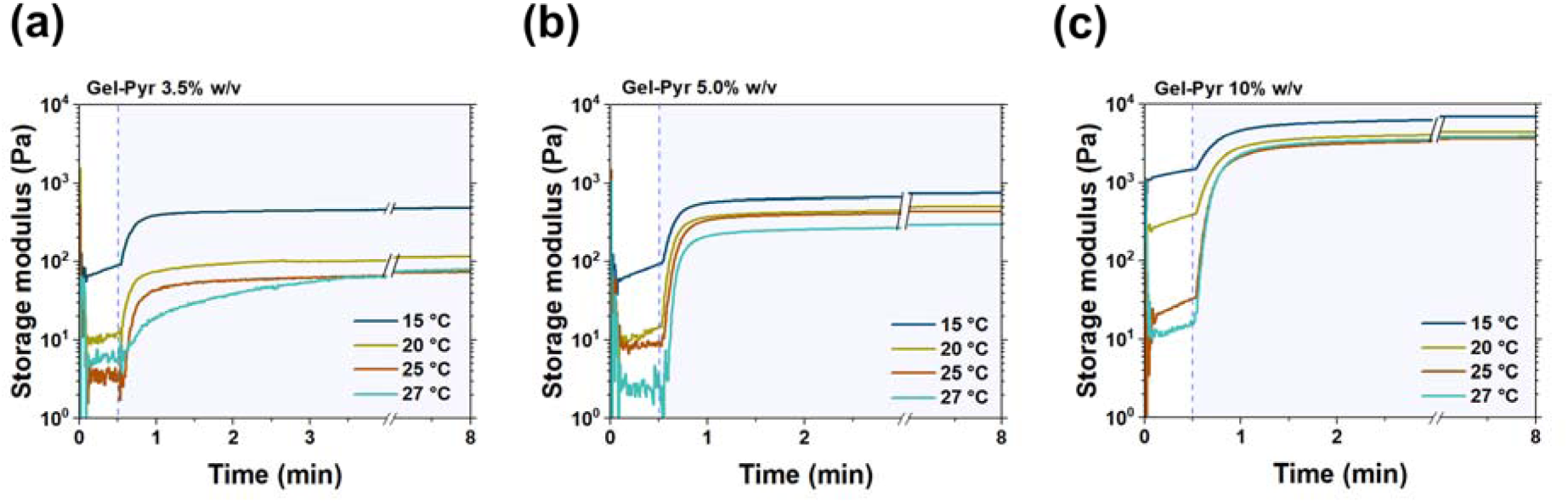
Photo-crosslinking evolution for Gel-Pyr at concentrations of (a) 3.5% w/v, (b) 5.0% w/v and (c) 10.0% w/v at a range of temperature from 15 to 27°C (I = 20 mW·cm^-^²).

### Compression testing

Figure S4 illustrates the stress-strain curves of Gel-Pyr at different concentrations. The compressive modulus was determined by applying a linear fit to the 10– 20% strain region of the curve. As shown, stress and strain exhibit a linear relationship at low strain levels (<30%), and all samples withstand the compression up to 80% strain.

**Fig. S4.**
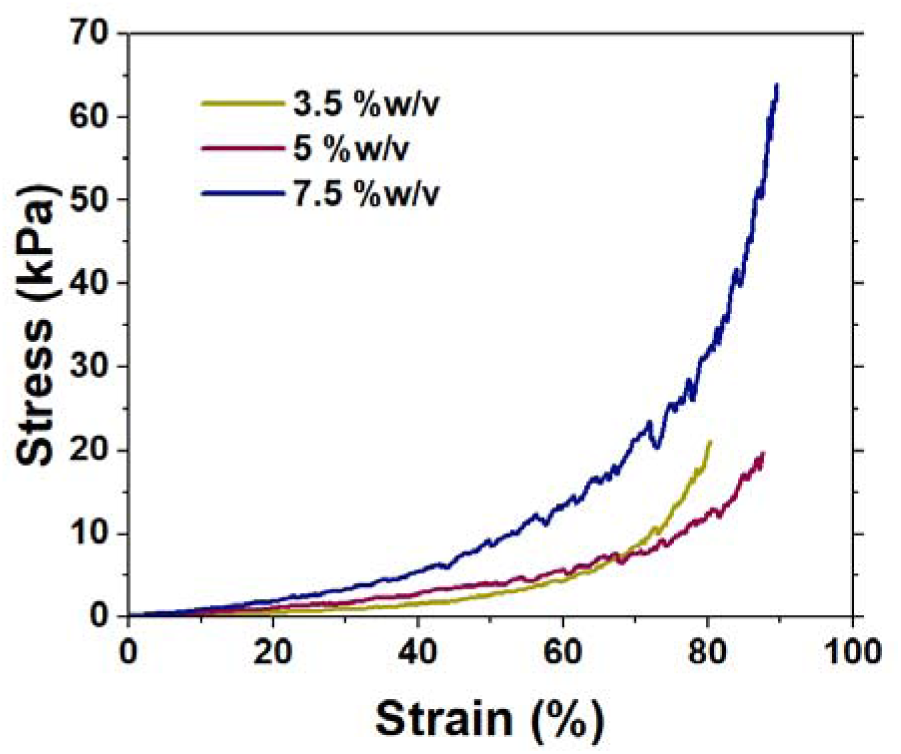
Stress-strain curve of Gel-Pyr at 3.5%, 5.0% and 7.5% w/v concentration.

### Flow behaviour as a function of temperature

The flow behaviour of 7.5% w/v Gel-Pyr was analysed using rotational rheology over a temperature range of 15 °C to 30 °C. Under nearly all tested conditions, Gel-Pyr exhibited shear-thinning behaviour (Figure S5). These findings were further utilized to estimate the shear stress and shear rate that the hydrogel precursors would experience under different bioprinting conditions.

**Fig. S5.**
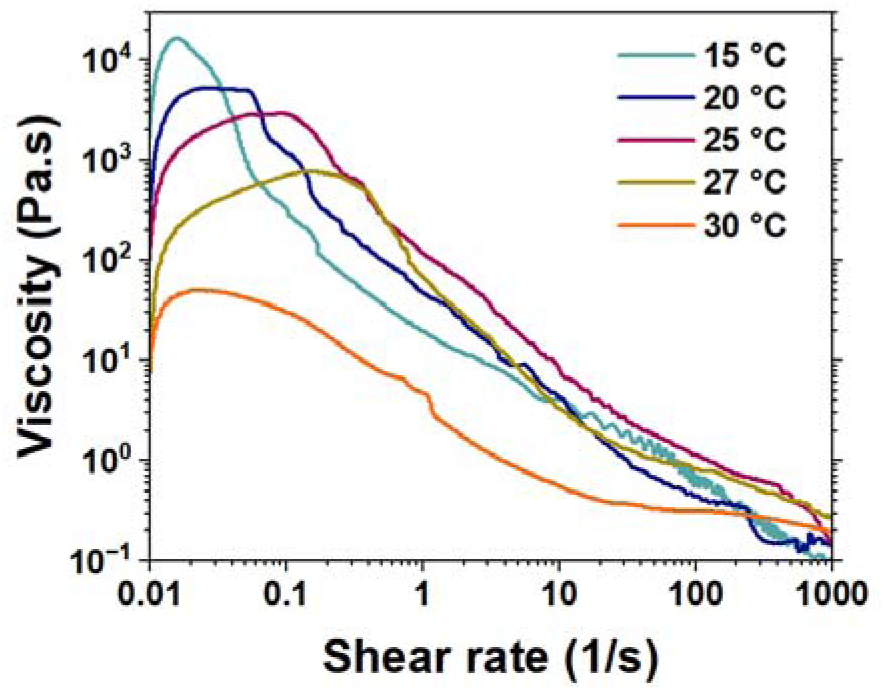
Viscosity under shear rate sweep of Gel-Pyr 7.5% w/v at a range of temperature from 15 to 30 °C.

**Table S1.** Consistency index (K) and Power-Law coefficient (n) for non-irradiated Gel-Pyr precursor 7.5% w/v at a range of temperature from 20 to 30 °C, where K and n are derived from linear regression of viscosity-shear rate profile (between 1 and 1000 s-1 for temperatures 20, 25 and 27°C; between 0.1 and 100 s-1 for temperature 30°C).

| Temperature (°C) | $n$ | $K$ |
| --- | --- | --- |
| 20 | 0.094 | 35.32 |
| 25 | 0.1 | 82.22 |
| 27 | 0.25 | 29.58 |
| 30 | 0.26 | 4.22 |

### Photo-rheology of GelMA with varying polymer and photoinitiator concentrations

To identify a GelMA formulation with mechanical properties comparable to Gel-Pyr, in situ photo-rheological analyses were performed using GelMA and lithium phenyl-2,4,6-trimethylbenzoylphosphinate (LAP) at varying concentrations under the same irradiation conditions as Gel-Pyr (λ = 400–500 nm, I = 20 mW·cm^-^², 25 °C). As shown in Figure S6, GelMA at 6% (w/v) exhibited a storage modulus comparable to Gel-Pyr (7.5% w/v) when prepared with either 0.5% or 1% (w/v) LAP. Therefore, 0.5% (w/v) LAP was selected for subsequent experiments to minimise photoinitiator concentration while maintaining comparable mechanical properties.

**Fig. S6.**
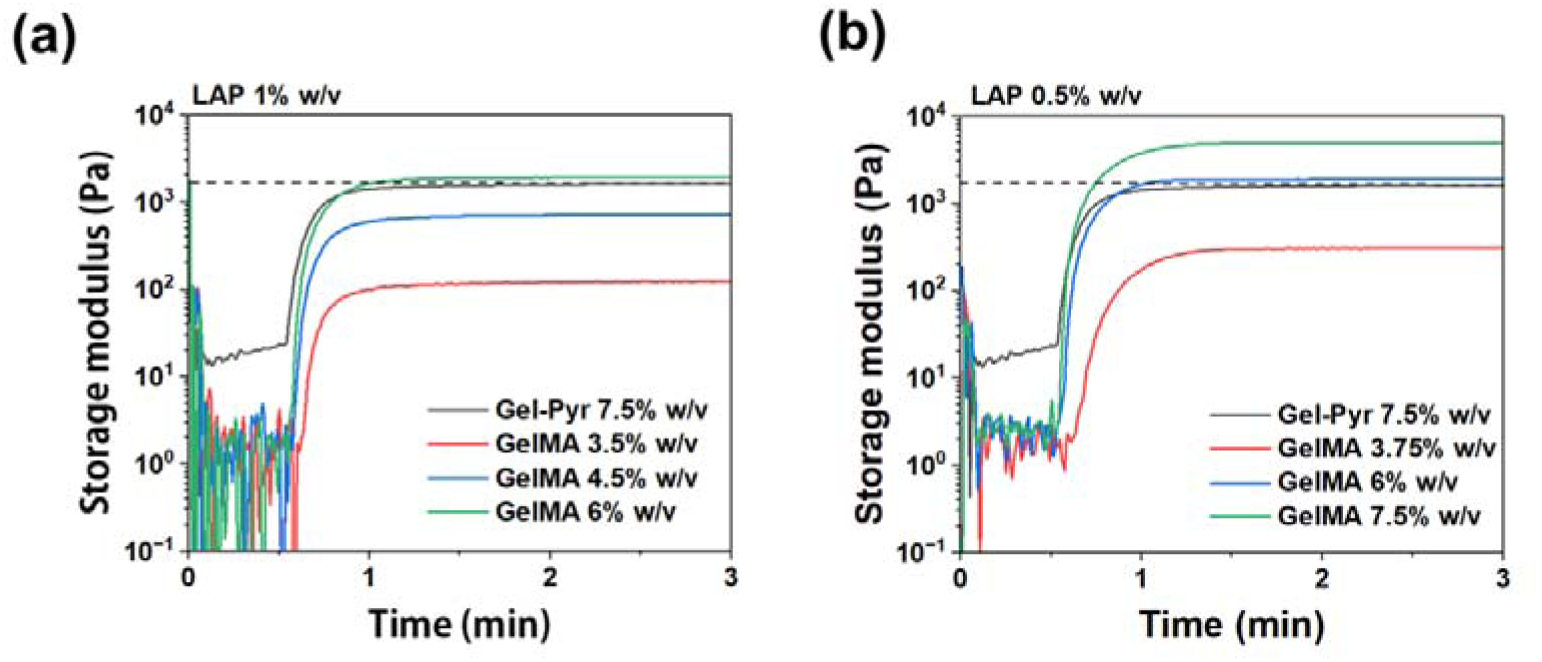
Photo-rheological analysis of GelMA hydrogels with varying polymer concentrations using (a) 1% w/v and (b) 0.5% w/v LAP.

### Mathematical model

The wall shear rate (γ̇) in the bioprinting nozzle is determined using the Metzner & Reed model (Eq. 1), while the shear stress (τ) (Eq. 2) and viscosity (η) (Eq. 3) are calculated based on the power-law model.

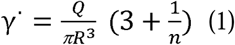

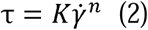

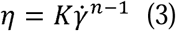

Where is the shear rate, *n* is the shear-thinning index, *Q* denotes volumetric flow rate, *R* is the nozzle radius, τ corresponds to shear stress, η is viscosity, and *K* is the consistency index. These equations use the rheological data (*n* and *K* derived from flow curves) and printing parameters (nozzle radius and extrusion speed) to estimate the shear stress and shear rate experienced during bioprinting. This predictive approach ensures that the mechanical stress imposed on the encapsulated cells remains within a suitable range, minimizing potential adverse effects on cell viability throughout the printing process.

## Notes

### Competing Interest Statement

The authors have declared no competing interest.

### Summary of Updates

The title and the discussion are updated for more clarity.

## References

1. Zhao Z, Vizetto-Duarte C, Moay ZK, Setyawati MI, Rakshit M, Kathawala MH, Ng KW. Composite hydrogels in three-dimensional in vitro models. Frontiers in Bioengineering and Biotechnology. 2020;8:611.

2. Jacob S, Nair AB, Shah J, Sreeharsha N, Gupta S, Shinu P. Emerging role of hydrogels in drug delivery systems, tissue engineering and wound management. Pharmaceutics. 2021;13(3):357.

3. Radhouani H, Correia S, Gonçalves C, Reis RL, Oliveira JM. Synthesis and characterization of biocompatible methacrylated kefiran hydrogels: Towards tissue engineering applications. Polymers. 2021;13(8):1342.

4. Kim J, Lee K-W, Hefferan TE, Currier BL, Yaszemski MJ, Lu L. Synthesis and evaluation of novel biodegradable hydrogels based on poly (ethylene glycol) and sebacic acid as tissue engineering scaffolds. Biomacromolecules. 2008;9(1):149–57.

5. Heinzmann C, Coulibaly S, Roulin A, Fiore GL, Weder C. Light-induced bonding and debonding with supramolecular adhesives. ACS applied materials & interfaces. 2014;6(7):4713–9.

6. Khayambashi P, Iyer J, Pillai S, Upadhyay A, Zhang Y, Tran SD. Hydrogel encapsulation of mesenchymal stem cells and their derived exosomes for tissue engineering. International journal of molecular sciences. 2021;22(2):684.

7. Rizwan M, Fokina A, Kivijärvi T, Ogawa M, Kufleitner M, Laselva O, et al. Photochemically activated Notch signaling hydrogel preferentially differentiates human derived hepatoblasts to cholangiocytes. Advanced Functional Materials. 2021;31(5):2006116.

8. Qazi TH, Blatchley MR, Davidson MD, Yavitt FM, Cooke ME, Anseth KS, Burdick JA. Programming hydrogels to probe spatiotemporal cell biology. Cell Stem Cell. 2022;29(5):678–91.

9. Yue K, Trujillo-de Santiago G, Alvarez MM, Tamayol A, Annabi N, Khademhosseini A. Synthesis, properties, and biomedical applications of gelatin methacryloyl (GelMA) hydrogels. Biomaterials. 2015;73:254–71.

10. Spearman BS, Agrawal NK, Rubiano A, Simmons CS, Mobini S, Schmidt CE. Tunable methacrylated hyaluronic acid based hydrogels as scaffolds for soft tissue engineering applications. Journal of Biomedical Materials Research Part A. 2020;108(2):279–91.

11. Hasany M, Talebian S, Sadat S, Ranjbar N, Mehrali M, Wallace GG, Mehrali M. Synthesis, properties, and biomedical applications of alginate methacrylate (ALMA)-based hydrogels: Current advances and challenges. Applied Materials Today. 2021;24:101150.

12. Mironi-Harpaz I, Wang DY, Venkatraman S, Seliktar D. Photopolymerization of cell-encapsulating hydrogels: crosslinking efficiency versus cytotoxicity. Acta biomaterialia. 2012;8(5):1838–48.

13. Nichol JW, Koshy ST, Bae H, Hwang CM, Yamanlar S, Khademhosseini A. Cell-laden microengineered gelatin methacrylate hydrogels. Biomaterials. 2010;31(21):5536–44.

14. Bowman CN, Kloxin CJ. Toward an enhanced understanding and implementation of photopolymerization reactions. AIChE Journal. 2008;54(11):2775–95.

15. Elkhoury K, Zuazola J, Vijayavenkataraman S. Bioprinting the future using light: A review on photocrosslinking reactions, photoreactive groups, and photoinitiators. SLAS technology. 2023;28(3):142–51.

16. Fairbanks BD, Schwartz MP, Bowman CN, Anseth KS. Photoinitiated polymerization of PEG-diacrylate with lithium phenyl-2, 4, 6-trimethylbenzoylphosphinate: polymerization rate and cytocompatibility. Biomaterials. 2009;30(35):6702-7.

17. Williams CG, Malik AN, Kim TK, Manson PN, Elisseeff JH. Variable cytocompatibility of six cell lines with photoinitiators used for polymerizing hydrogels and cell encapsulation. Biomaterials. 2005;26(11):1211–8.

18. Xu L, Sheybani N, Yeudall WA, Yang H. The effect of photoinitiators on intracellular AKT signaling pathway in tissue engineering application. Biomaterials science. 2015;3(2):250–5.

19. Urushibara A, Kodama S, Yokoya A. Induction of genetic instability by transfer of a UV-A-irradiated chromosome. Mutation Research/Genetic Toxicology and Environmental Mutagenesis. 2014;766:29–34.

20. Asim S, Tuftee C, Qureshi AT, Callaghan R, Geary ML, Santra M, et al. Multi Functional Gelatin Dithiolane Hydrogels for Tissue Engineering. Advanced Functional Materials. 2025;35(3):2407522.

21. Velayudhan S, Pr AK. Biocompatibility evaluation of antioxidant cocktail loaded gelatin methacrylamide as bioink for extrusion-based 3D bioprinting. Biomedical Materials. 2023;18(4):044101.

22. Pai RR, Ajit S, Nair SS, Kumar PA, Velayudhan S. Radical scavenging gelatin methacrylamide based bioink formulation for three dimensional bioprinting of parenchymal liver construct. Bioprinting. 2022;27:e00214.

23. Dogan E, Austin A, Pourmostafa A, Yogeshwaran S, Hosseinabadi HG, Miri AK. Design considerations for photoinitiator selection in cell-laden gelatin methacryloyl hydrogels. Biomaterials science. 2026;14(3):807–16.

24. Truong VX, Li F, Ercole F, Forsythe JS. Wavelength-selective coupling and decoupling of polymer chains via reversible [2+ 2] photocycloaddition of styrylpyrene for construction of cytocompatible photodynamic hydrogels. ACS Macro Letters. 2018;7(4):464–9.

25. Koay WL, Gao C, Vu QT, Oh XY, Lin H, Mondal S, et al. Light Switchable Bioorthogonal Reaction Manifold for Modulation of Hydrogel Properties. Biomacromolecules. 2024;25(10):6635–44.

26. Picchiotti A, Nenov A, Giussani A, Prokhorenko VI, Miller RD, Mukamel S, Garavelli M. Pyrene, a test case for deep-ultraviolet molecular photophysics. The Journal of Physical Chemistry Letters. 2019;10(12):3481–7.

27. Hafidz R, Yaakob C, Amin I, Noorfaizan A. Chemical and functional properties of bovine and porcine skin gelatin. International Food Research Journal. 2011;18(2):787–91.

28. Hermanto S, Fatimah W. Differentiation of bovine and porcine gelatin based on spectroscopic and electrophoretic analysis. Journal of Food and Pharmaceutical Sciences. 2013;1(3).

29. Pepelanova I, Kruppa K, Scheper T, Lavrentieva A. Gelatin-methacryloyl (GelMA) hydrogels with defined degree of functionalization as a versatile toolkit for 3D cell culture and extrusion bioprinting. Bioengineering. 2018;5(3):55.

30. Li X, Chen S, Li J, Wang X, Zhang J, Kawazoe N, Chen G. 3D culture of chondrocytes in gelatin hydrogels with different stiffness. Polymers (Basel). 2016; 8 (8). Epub 2016/07/26. doi: 10.3390/polym8080269. PubMed PMID: 30974547.

31. Stojkov G, Niyazov Z, Picchioni F, Bose RK. Relationship between structure and rheology of hydrogels for various applications. Gels. 2021;7(4):255.

32. Alam K, Iqbal M, Hasan A, Al-Maskari N. Rheological characterization of biological hydrogels in aqueous state. Journal of Applied Biotechnology Reports. 2020;7(3):171–5.

33. Silva Correia J, Gloria A, Oliveira MB, Mano JF, Oliveira JM, Ambrosio L, Reis RL. Rheological and mechanical properties of acellular and cell laden methacrylated gellan gum hydrogels. Journal of Biomedical Materials Research Part A: An Official Journal of The Society for Biomaterials, The Japanese Society for Biomaterials, and The Australian Society for Biomaterials and the Korean Society for Biomaterials. 2013;101(12):3438–46.

34. Yao Y, Molotnikov A, Parkington HC, Meagher L, Forsythe JS. Extrusion 3D bioprinting of functional self-supporting neural constructs using a photoclickable gelatin bioink. Biofabrication. 2022;14(3):035014.

35. Lee JH, Bucknall DG. Swelling behavior and network structure of hydrogels synthesized using controlled UV initiated free radical polymerization. Journal of Polymer Science Part B: Polymer Physics. 2008;46(14):1450–62.

36. Gładysiak A, Nguyen TN, Bounds R, Zacharia A, Itskos G, Reimer JA, Stylianou KC. Temperature-dependent interchromophoric interaction in a fluorescent pyrene-based metal– organic framework. Chemical science. 2019;10(24):6140–8.

37. Pise S, Chatterjee A, Dey N. Exploring Self Assembly and Viscoelastic Behavior of Pyrene Based Fluorescent Hydrogel: Designing Paper Sensors for Water Soluble Explosives. European Journal of Organic Chemistry. e202401096.

38. Fiorica C, Biscari G, Palumbo FS, Pitarresi G, Martorana A, Giammona G. Physicochemical and rheological characterization of different low molecular weight gellan gum products and derived ionotropic crosslinked hydrogels. Gels. 2021;7(2):62.

39. Steyaert I, Rahier H, Van Vlierberghe S, Olijve J, De Clerck K. Gelatin nanofibers: Analysis of triple helix dissociation temperature and cold-water-solubility. Food Hydrocolloids. 2016;57:200–8.

40. Joly-Duhamel C, Hellio D, Djabourov M. All gelatin networks: 1. Biodiversity and physical chemistry. Langmuir. 2002;18(19):7208–17.

41. Michon C, Cuvelier G, Relkin P, Launay B. Influence of thermal history on the stability of gelatin gels. International Journal of Biological Macromolecules. 1997;20(4):259–64.

42. Kokol V, Pottathara YB, Mihelčič M, Perše LS. Rheological properties of gelatine hydrogels affected by flow-and horizontally-induced cooling rates during 3D cryo-printing. Colloids and Surfaces A: Physicochemical and Engineering Aspects. 2021;616:126356.

43. Ha S, Lee Y, Kwak Y, Mishra A, Yu E, Ryou B, Park C-M. Alkyne–Alkene [2+ 2] cycloaddition based on visible light photocatalysis. Nature Communications. 2020;11(1):2509.

44. Podborska A, Suchecki M, Mech K, Marzec M, Pilarczyk K, Szaciłowski K. Light intensity-induced photocurrent switching effect. Nature Communications. 2020;11(1):854.

45. Guvendiren M, Lu HD, Burdick JA. Shear-thinning hydrogels for biomedical applications. Soft matter. 2012;8(2):260–72.

46. Ali I, Shah LA. Rheological investigation of the viscoelastic thixotropic behavior of synthesized polyethylene glycol-modified polyacrylamide hydrogels using different accelerators. Polymer Bulletin. 2021;78(3):1275–91.

47. Wang Q, Karadas Ö, Rosenholm JM, Xu C, Näreoja T, Wang X. Bioprinting macroporous hydrogel with aqueous two phase emulsion based bioink: in vitro mineralization and differentiation empowered by phosphorylated cellulose nanofibrils. Advanced Functional Materials. 2024;34(29):2400431.

48. Rastin H, Ormsby RT, Atkins GJ, Losic D. 3D bioprinting of methylcellulose/gelatin-methacryloyl (MC/GelMA) bioink with high shape integrity. ACS Applied Bio Materials. 2020;3(3):1815–26.

49. Müller SJ, Fabry B, Gekle S. Predicting cell stress and strain during extrusion bioprinting. Physical Review Applied. 2023;19(6):064061.

50. Ouyang L, Yao R, Zhao Y, Sun W. Effect of bioink properties on printability and cell viability for 3D bioplotting of embryonic stem cells. Biofabrication. 2016;8(3):035020.

51. Hendriks J, Willem Visser C, Henke S, Leijten J, Saris DB, Sun C, et al. Optimizing cell viability in droplet-based cell deposition. Scientific reports. 2015;5(1):11304.

52. Lopez Hernandez H, Souza JW, Appel EA. A quantitative description for designing the extrudability of shear thinning physical hydrogels. Macromolecular Bioscience. 2021;21(2):2000295.

53. Schwab A, Levato R, D’Este M, Piluso S, Eglin D, Malda J. Printability and shape fidelity of bioinks in 3D bioprinting. Chemical reviews. 2020;120(19):11028–55.

54. Paxton N, Smolan W, Böck T, Melchels F, Groll J, Jungst T. Proposal to assess printability of bioinks for extrusion-based bioprinting and evaluation of rheological properties governing bioprintability. Biofabrication. 2017;9(4):044107.

55. Wang L, Xu M-e, Luo L, Zhou Y, Si P. Iterative feedback bio-printing-derived cell-laden hydrogel scaffolds with optimal geometrical fidelity and cellular controllability. Scientific reports. 2018;8(1):2802.

56. Li M, Tian X, Zhu N, Schreyer DJ, Chen X. Modeling process-induced cell damage in the biodispensing process. Tissue Engineering Part C: Methods. 2010;16(3):533–42.

57. Lee GM, Kim S-j, Kim EM, Kim E, Lee S, Lee E, et al. Free radical-scavenging composite gelatin methacryloyl hydrogels for cell encapsulation. Acta Biomaterialia. 2022;149:96–110.

58. Ansari A, Bhowmik S, Zhang K, Reeves CL, Vahala D, Choi YS, et al. A Visible Light Responsive Hydrogel to Study the Effect of Dynamic Tissue Stiffness on Cellular Mechanosensing. Advanced Functional Materials. 2025;35(35):2501585.

59. Zhang Y, Lv J, Zhao J, Ling G, Zhang P. A versatile GelMA composite hydrogel: designing principles, delivery forms and biomedical applications. European Polymer Journal. 2023;197:112370.

60. Paul S, Schrobback K, Tran PA, Meinert C, Davern JW, Weekes A, Klein TJ. Photo cross linkable, injectable, and highly adhesive GelMA glycol chitosan hydrogels for cartilage repair. Advanced Healthcare Materials. 2023;12(32):2302078.

61. Das S, Jegadeesan JT, Basu B. Gelatin methacryloyl (GelMA)-based biomaterial inks: process science for 3D/4D printing and current status. Biomacromolecules. 2024;25(4):2156–221.

62. Hou J, Han Z, Jing Y, Yang X, Zhang S, Sun K, et al. Autophagy prevents irradiation injury and maintains stemness through decreasing ROS generation in mesenchymal stem cells. Cell death & disease. 2013;4(10):e844-e.

63. Borodkina A, Shatrova A, Abushik P, Nikolsky N, Burova E. Interaction between ROS dependent DNA damage, mitochondria and p38 MAPK underlies senescence of human adult stem cells. Aging (Albany NY). 2014;6(6):481.

64. Bigarella CL, Liang R, Ghaffari S. Stem cells and the impact of ROS signaling. Development. 2014;141(22):4206–18.

65. Zhao J, Zhang L, Lu A, Han Y, Colangelo D, Bukata C, et al. ATM is a key driver of NF-κB-dependent DNA-damage-induced senescence, stem cell dysfunction and aging. Aging (Albany NY). 2020;12(6):4688.

66. Lin CH, Li NT, Cheng HS, Yen ML. Oxidative stress induces imbalance of adipogenic/osteoblastic lineage commitment in mesenchymal stem cells through decreasing SIRT1 functions. Journal of cellular and molecular medicine. 2018;22(2):786–96.

67. Atashi F, Modarressi A, Pepper MS. The role of reactive oxygen species in mesenchymal stem cell adipogenic and osteogenic differentiation: a review. Stem cells and development. 2015;24(10):1150–63.

68. Denu RA, Hematti P. Optimization of oxidative stress for mesenchymal stromal/stem cell engraftment, function and longevity. Free Radical Biology and Medicine. 2021;167:193–200.

69. Yu K-R, Lee JY, Kim H-S, Hong I-S, Choi SW, Seo Y, et al. A p38 MAPK-mediated alteration of COX-2/PGE2 regulates immunomodulatory properties in human mesenchymal stem cell aging. PloS one. 2014;9(8):e102426.

70. Stavely R, Nurgali K. The emerging antioxidant paradigm of mesenchymal stem cell therapy. Stem Cells Translational Medicine. 2020;9(9):985–1006.

